# Marine gregarine genomes reveal the breadth of apicomplexan diversity and provide new insights on gliding motility

**DOI:** 10.1101/2021.08.09.454911

**Authors:** Julie Boisard, Evelyne Duvernois-Berthet, Linda Duval, Joseph Schrével, Laure Guillou, Amandine Labat, Sophie Le Panse, Gérard Prensier, Loïc Ponger, Isabelle Florent

**Affiliations:** Molécules de Communication et Adaptation des Microorganismes (MCAM, UMR 7245 CNRS), Département Adaptations du vivant (AVIV), Muséum National d’Histoire Naturelle, CNRS, CP 52, 57 rue Cuvier, 75231 Paris Cedex 05, France; Structure et instabilité des génomes (STRING UMR 7196 CNRS/INSERM U1154), Département Adaptations du vivant (AVIV), Muséum National d’Histoire Naturelle, CNRS, INSERM, CP 26, 57 rue Cuvier, 75231 Paris Cedex 05, France; Department of Biology, Lund University, Sölvegatan 35, 223 62 Lund, Sweden; Muséum National d’Histoire Naturelle, Centre National de la Recherche Scientifique, Laboratoire Physiologie Moléculaire et Adaptation (PhyMA), UMR7221 CNRS-MNHN, 75005, Paris, France; Sorbonne Université, CNRS, UMR7144 Adaptation et Diversité en Milieu Marin, Ecology of Marine Plankton (ECOMAP), Station Biologique de Roscoff SBR, 29680 Roscoff, France; Plateforme d’Imagerie Merimage, FR2424, Centre National de la Recherche Scientifique, Station Biologique de Roscoff, Roscoff, 29680, France; Cell biology and Electron Microscopy Laboratory, François Rabelais University, 10 Boulevard Tonnellé, BP 3223 Tours Cedex, France

**Keywords:** Apicomplexa, marine gregarine, genome assembly, comparative genomics, gliding, phylogeny

## Abstract

Our current view of the evolutionary history, coding and adaptive capacities of Apicomplexa, protozoan parasites of a wide range of metazoan, is currently strongly biased toward species infecting humans, as data on early diverging apicomplexan lineages infecting invertebrates is extremely limited. Here, we characterized the genome of the marine eugregarine *Porospora gigantea*, intestinal parasite of Lobsters, remarkable for the macroscopic size of its vegetative feeding forms (trophozoites) and its gliding speed, the fastest so far recorded for Apicomplexa. Two highly syntenic genomes named A and B were assembled. Similar in size (~9 Mb) and coding capacity (~5300 genes), A and B genomes are 10.8% divergent at the nucleotide level, corresponding to 16-38 My in divergent time. Orthogroup analysis across 25 (proto)Apicomplexa species, including *Gregarina niphandrodes*, showed that A and B are highly divergent from all other known apicomplexan species, revealing an unexpected breadth of diversity. Phylogenetically these two species branch sister to Cephaloidophoroidea, and thus expand the known crustacean gregarine superfamily. The genomes were mined for genes encoding proteins necessary for gliding, a key feature of apicomplexans parasites, currently studied through the molecular model called glideosome. Sequence analysis shows that actin-related proteins and regulatory factors are strongly conserved within apicomplexans. In contrast, the predicted protein sequences of core glideosome proteins and adhesion proteins are highly variable among apicomplexan lineages, especially in gregarines. These results confirm the importance of studying gregarines to widen our biological and evolutionary view of apicomplexan species diversity, and to deepen our understanding of the molecular bases of key functions enabling parasitism, such as the glideosome.

## BACKGROUND

Apicomplexans are unicellular eukaryotic microorganisms that have evolved towards endobiotic symbionts or parasites. The Apicomplexa include about 350 genera^1^ for 6,000 documented species. Some species are extremely pathogenic such as *Plasmodium* spp., *Toxoplasma gondii* and *Cryptosporidium* spp., responsible for malaria, toxoplasmosis and cryptosporidiosis, respectively. Current knowledge of apicomplexan genomes is based on sequence data from a dozen genera, and more precisely, the genera which include highly pathogenic species^2^. Consequently, our view of the Apicomplexa genome is highly skewed towards intracellular parasites of vertebrates, notably Coccidia, Hemosporidia and *Cryptosporidium* (see references in Table S1). By comparison, the gregarines, of which there are at least 1,770 species^3^, have hardly been explored at an omic level^4^. Gregarines were identified as the most abundant and widely reported apicomplexan in a recent environmental study^5^. However, as they have low pathogenicity and are non-cultivable in the laboratory, they have attracted less interest.

Overlooking the gregarines risks leaving part of the evolutionary history of Apicomplexa unexplored, because they represent early diverging lineages as well as displaying a diversity of specific adaptive traits. For instance, gregarines are mostly extracellular, infecting a wide diversity of marine and terrestrial non-vertebrate hosts^6,7^. At this time, available genomic data are very limited to terrestrial gregarines, such as partial data on *Ascogregarina taiwanensis*, an intestinal parasite of the tiger mosquito *Aedes albopictus*^8^, and the draft genome of *Gregarina niphandrodes*, an intestinal parasite of the mealworm *Tenebrio molitor* (unpublished, available in CryptoDB^9^). Transcriptomic studies on trophozoite (feeding) stages of terrestrial and marine gregarine species have recently provided important insights^10–13^, especially about organellar genomes and metabolic pathways. These developmental stage-dependent data, however, do not provide a complete picture of the genetic landscape of gregarines, nor can they provide information on their genome structure.

To study the gregarine genome, we focused on the marine eugregarine *Porospora gigantea* (Van Beneden, 1869) Schneider, 1875, which is an intestinal parasite of the lobster *Homarus gammarus*. First described in 1869, E. Van Beneden named the organism *Gregarina gigantea* in reference to the “gigantic” size (up to 16,000 μm) of the trophozoite stages, being visible to the naked eye^14^. Van Beneden reported that “cyst” forms of this parasite accumulated within the chitinous folds of the lobster rectum, the “rectal ampulla”. Schneider went on to show that these cysts enclosed thousands of “gymnospores” or “heliospores”, corresponding to spherical groups of very tiny zoites radiating from a central, optically void mass, and renamed the species *Porospora gigantea* (van Beneden, 1869) Schneider, 1875^15^. Biological material for genomic studies is particularly difficult to gather from non-cultivable microorganisms, so we took advantage of the existence of these well-described structures^16–19^, knowing that each cyst contains several thousand “gymnospores”, each composed of hundreds of zoites, involving the natural amplification of its genomic material. Cysts indeed proved to be a remarkable natural source of genomic DNA. Gliding is a characteristic apicomplexan movement that also happens to be essential for the invasion and egress of host cells, and thus for the intracellular parasitic lifestyle^20–24^. *P. gigantea* trophozoites are known to glide at rates of up to 60μm/s^25^, so are prime candidates in which to study the mechanism of gliding motility. Currently about 40 proteins, identified mainly in *T. gondii* and *Plasmodium falciparum*, compose the glideosome, a commonly accepted structural model of the apicomplexan motor complex (see Frénal et al, 2017^26^ for review).

In this study, we report the assembly of the first two draft genomes of *P. gigantea*. We present their main features and predicted proteomes and compare them to other available apicomplexan genomes, revealing an unexpected diversity. We investigated their position within Apicomplexa and among the major subgroups of gregarines through a phylogenomic analysis. We also examined their position within the crustacean gregarines according to 18S ribosomal gene sequences. Finally, a comparative study was performed to gain insight into the conservation of gliding proteins for these gregarines, the currently fastest moving extracellular Apicomplexa.

## RESULTS

### Phenotypic characterization

Specimens of the lobster *Homarus gammarus*, the type host species for *Porospora gigantea*, were collected either from the sea in Roscoff bay (France) or from commercial lobster tanks in Roscoff (Figure 1, Table S2). A total of 35 lobsters (9 from the wild and 26 from captivity) were dissected and infection with *P. gigantea* was quantified (Figure 1, Figure S1). Overall, infection levels were significantly higher in lobsters freshly caught from the sea (prevalence of 100%, high parasitic loads) than in lobsters that had been held in captivity in lobster tanks (prevalence < 62%, low parasitic loads, see Table S2), a similar result to that reported by Van Beneden (1869)^14^. The morphology of cysts, gymnospores, zoites and trophozoites was imaged and measured (Figure 1, Tables S3, S4 and S5). Cysts were mostly spherical but some were ovoid, with diameters ranging from ~108μm to ~240μm (mean ± standard deviation, 151.1 ± 45.3μm, n = 97), and they enclosed thousands of gymnospores, that were also mostly spherical, with diameters from less than 5μm to almost 7μm (5.63 ± 0.69μm, n = 265). These gymnospores were indeed composed of radially arranged zoites forming a monolayer with an optically void center. Observation of broken gymnospores by scanning electron microscopy made it possible to measure the length of the constituent zoites (1.04 ± 0.16μm, n = 105) and their apical width (0.630 ± 0.129μm, n = 176). Trophozoites were very thin and long, up to 2585μm for a mean width of 41.8 ± 10.4μm (n = 104). As previously described, the posterior of the trophozoite was slightly thinner, ~30μm. The whole trophozoite surface was covered by longitudinal epicytic folds (Figure S1.B) that are thought to be necessary for eugregarine gliding^27^. The sum of these morphological observations all accord with the species being *P. gigantea* from the type host *H. gammarus*^6,14,15^.

**Figure 1.**
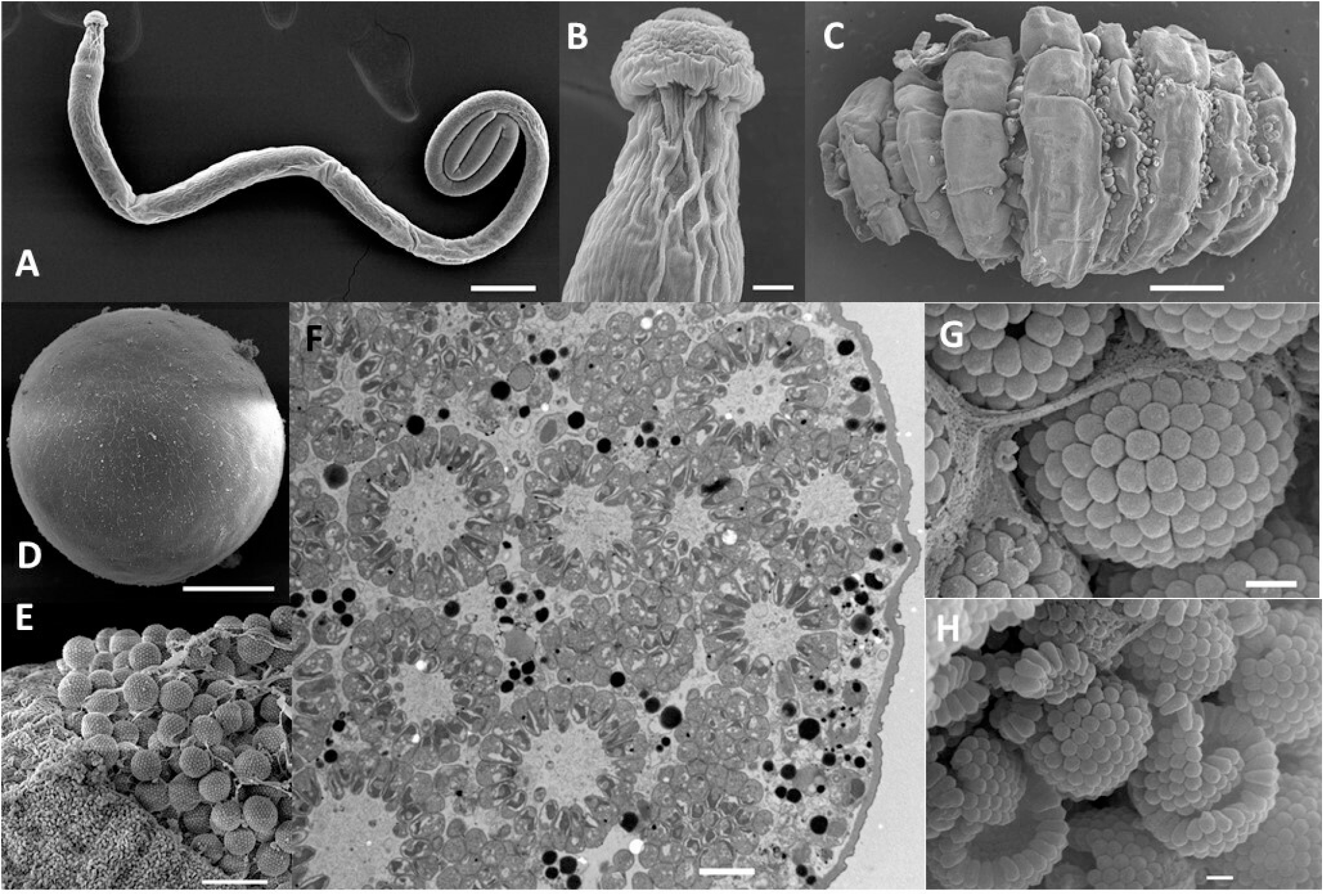
Morphological characterization of *Porospora cf. gigantea*. **A.** Trophozoite stage (Tropho #8, Lobster #12) (scale bar = 100μm). **B.** Zoom on A, showing trophozoite epimerite (scale bar = 10μm). **C**. Rectal ampulla showing cysts in folds (Lobster #4) (scale bar = 1 mm). **D.** Isolated cyst (Cyst #4, Lobster #12) (scale bar = 50μm). **E.** Broken cyst packed with gymnospores (Lobster #4) (scale = 10 μm). **F**. Section across a cyst illustrating radial arrangement of zoites in gymnospores (JS449 = Lobster #35) (scale bar = 2μm). **G., H.** Zoom on intact and broken gymnospores showing zoites (Lobster #4) (scale = 1μm). All images are scanning electronic micrographs except F which is a transmission electronic micrograph. See also Figure S1, Tables S2, S3, S4, S5 and S6.

Gliding of isolated trophozoites was filmed. The dynamic recordings confirm that trophozoites moved uni-directionally, with the protomerite forwards, in either straight or curved lines depending on the individuals observed, with the whole body (deutomerite) following the same path as the apical protomerite (Film S1). The speed of trophozoite displacement was estimated to be ~60μm/sec, as initially observed by King and Sleep (2005)^25^, but was faster than 100μm/sec in some recordings (Table S6). No syzygy was observed. Some solitary encysted trophozoites were observed, supporting the observation of Léger and Duboscq (1909)^28^, who considered that encysted gymnospores correspond to a schizogonic rather than a gamogonic phase of *Porospora* development, a hypothesis that is still debated^6^.

### Two highly related genomes

Four biological samples were sequenced and analyzed independently, and then assembled together (Figure 2.A). The raw assembly produced 214,938 contigs (99.6 Mb) among which were 13,656 contigs longer than 1 kb (47.9 Mb). The scaffolds obtained were cleaned by removing contaminants such as bacterial, fungal and host sequences (Figure 2.B), resulting in a raw assembly of 1719 contigs covering 18 Mb.

**Figure 2.**
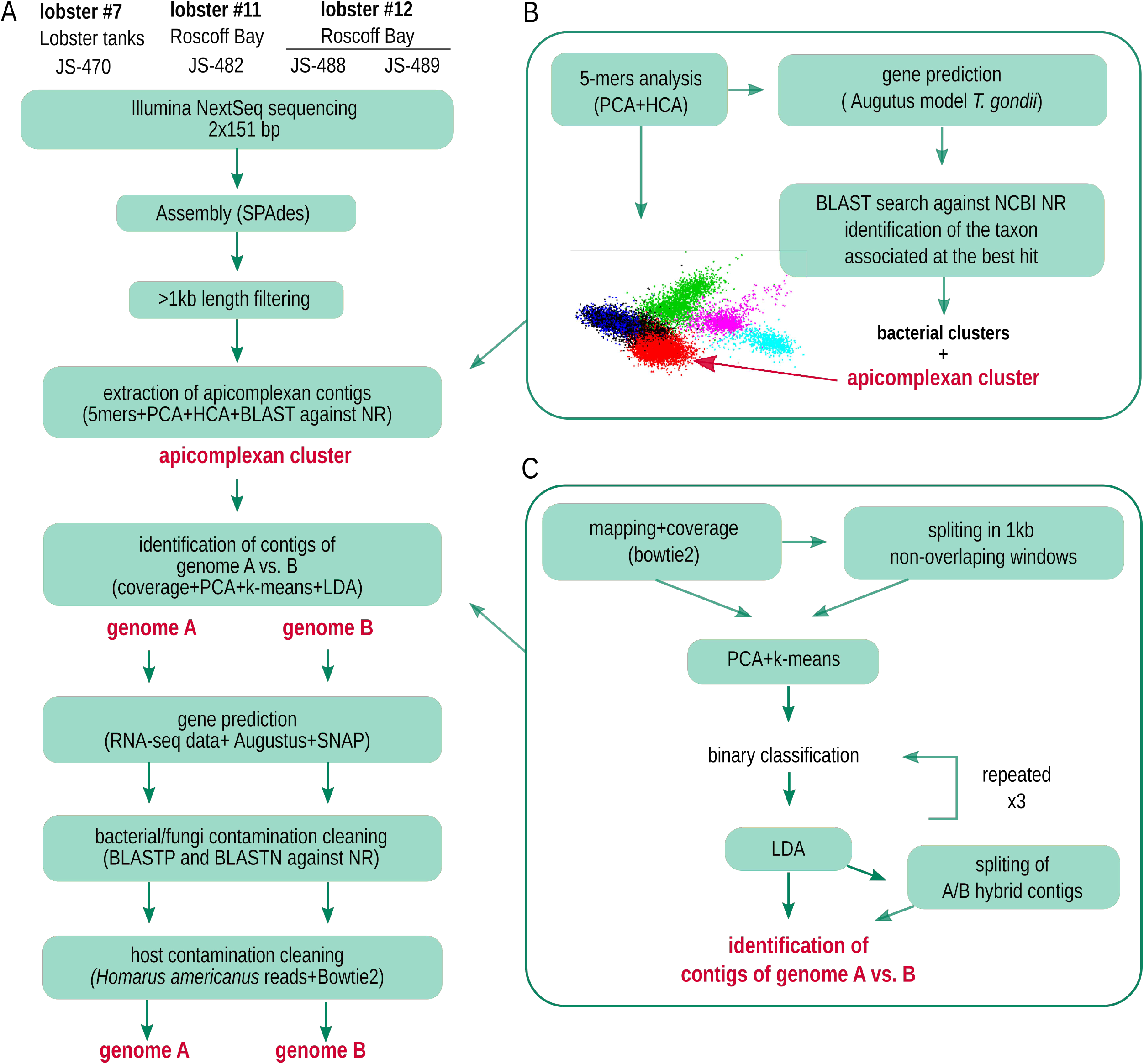
Protocol for assembling the two genomes. **A.** Overview of the full protocol. **B.** Identification of apicomplexan vs contaminant contigs based on k-mer composition. **C.** Identification of contigs from genomes A and B based on coverage data for each individual library. See also Figures S2, S3 and S4.

The analysis of contig coverage for each individual library revealed a bimodal distribution suggesting a mixture of genomes in differing proportions depending on the biological sample (Figure S2). More precisely, while only one set of contigs displayed a significant coverage for the lobster tank parasite sample (JS-470, peak around 250×), the three other parasite samples from freshly captured hosts (JS-482, JS-488, JS-489) showed two distinct sets of scaffolds with similar size (~9Mb) and different coverage values. The difference in coverage was used to split the whole assembled contigs into two sets that were named A for the set of contigs present in all four samples, and B for the set present only in the three lobsters freshly captured in the wild (Figure 2.C). The percentages of genomes A and B in each biological DNA sample was estimated (Figure S2) as 100% A for JS-470, 63.2% A and 36.8% B for JS-482, 70.5% A and 29.5% B for JS-488, and 62.4% A and 37.6% B for JS-489, based on medium coverage levels. Genome A maps to 787 contigs for a total of 8.8 Mb, whereas genome B maps to 933 contigs for a total of 9.0 Mb. Contigs from the two genomes can be aligned with each other over 7.7Mb, with a percentage of divergence around 10.8% at the nucleotide level.

To summarize, the A and B genomes associated to the species named *P. cf. gigantea* are similar in size (~9Mb) and are syntenic but divergent (Figure 2).

### Genome features

#### Two genomes with similar coding capacities

A total of 10,631 putative genes were predicted from the raw assembly (17,930 including alternative splicing), which were split into two sets of similar size: 5270 genes in genome A (8835 transcripts) and 5361 genes (9035 transcripts) in genome B (Table 1, Figure 2). The completeness of both A and B genomes was assessed by using BUSCO software^29^ on the Apicomplexa geneset (n = 446). Genomes A and B respectively showed completeness scores of 70% (n = 312) and 67.7% (n = 302) (Figure S3).

**Table 1.** Metrics of the genomes of *P. cf. gigantea* and a selection of 6 reference species. See also Figure S1 and S2.

| species | *P. cf. gigantea* |  | *G. niphandrodes* | *C. parvum* | *T. gondii* | *P. falciparum* | *C. velia* | *V. brassicaformis* |
| --- | --- | --- | --- | --- | --- | --- | --- | --- |
| strain | A | B | na | lowall | ME49 | 3D7 | CCMP2878 | CCMP3155 |
| nb of contigs/chromosomes | 787 | 934 | 355 | 8 | 435 | 14 | 5470 | 1006 |
| total length of assembly (bp) | 8806768 | 9049943 | 13873624 | 9102324 | 63472444 | 23292622 | 192006978 | 72475329 |
| mean length contigs/chromosomes (bp) | 11190.3 | 9689.45 | 39080.63 | 1137790.5 | 145913.66 | 1663758.71 | 35101.82 | 72043.07 |
| GC content (%) | 54.3 | 54.3 | 53.8 | 30.2 | 52.4 | 19.3 | 49.1 | 58.1 |
| nb of protein coding genes | 5270 | 5361 | 6606 | 4020 | 8862 | 5602 | 30604 | 23412 |
| mean length of coding genes (bp) | 1438.2 | 1450.3 | 1392.6 | 1865.0 | 5602.9 | 2488.6 | 4507.6 | 2704.7 |
| nb of tRNA | 14 | 14 | 231 | 45 | 150 | 45 | 0 | 0 |
| nb of rRNA | 27 | 25 | 0 | 5 | 420 | 28 | 0 | 0 |
| nb of gene with intron(s) | 2957 | 2981 | 2390 | 575 | 6801 | 3010 | 21895 | 22163 |
| median length of the introns (bp) | 28<br>[27-30] | 28<br>[27-30] | 95<br>[56-145] | 65<br>[51-91] | 467<br>[322-632] | 140<br>[110-184] | 372<br>[273-520] | 81<br>[70-98] |
| mode of intron length (bp) | 28 | 28 | 37 | 44 | 55 | 121 | 320 | 74 |
| mean nb of introns per gene* | 1.8 | 1.8 | 1.4 | 1.8 | 5.9 | 2.9 | 5.4 | 7.9 |
| non-coding DNA (%) | 16 | 16 | 37 | 24 | 68 | 47 | 74 | 50 |

\* by considering only genes with intron(s)

The number of A and B orthologues was investigated. The predicted proteins of *P. cf. gigantea* A and B were split into 5656 orthogroups including 4443 groups (88%) which had at least one orthologous gene for both A and B. This percentage of common orthogroups between genomes A and B is higher than that observed between *Plasmodium falciparum* and *Plasmodium berghei* (70%), thought to have diverged around 33 Mya ago (TimeTree^30^), but similar to that observed between *P. falciparum* and *Plasmodium reichenowi* (86%, 3.3 – 7.7 Mya, TimeTree).

The percentages of shared orthogroups between *P. cf. gigantea* genomes and each of the reference apicomplexan species are similar (*Cryptosporidium parvum*, 18%; *G. niphandrodes*, 17%; *P. falciparum*, 14%; *T. gondii*, 14%) despite the differences in divergence, but it is higher than the percentages observed with chromerid species (*Chromera velia*, 8%; *Vitrella brassicaformis*, 10%). We can deduce from these results that the *P. cf. gigantea* genomes do not share significantly more orthogroups with *G. niphandrodes*, the only other available gregarine genome, than with any other apicomplexan (Figure 3).

**Figure 3.**
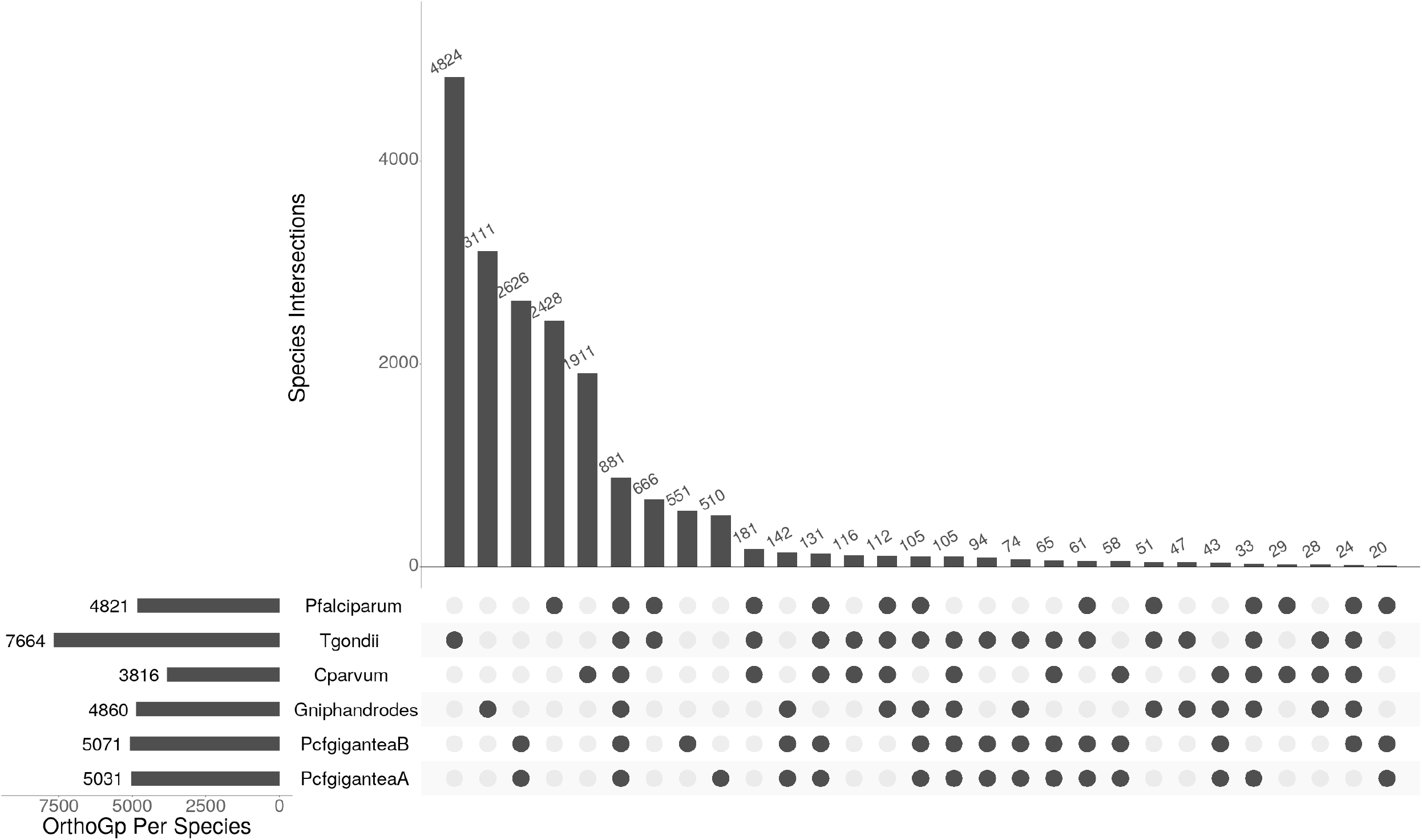
Shared apicomplexan proteins. Distribution of the orthogroups among *P. cf. gigantea* A and B and 4 species of apicomplexans: the gregarine *G. niphandrodes*, the cryptosporidian *C. parvum*, the coccidian *T. gondii* and the hematozoan *P. falciparum*. Only bars with more than 20 orthogroups are shown. See also Table S1.

#### Two gene-dense genomes with small introns

The proportion of coding sequences in A and B genomes is 84%, which is particularly high compared to other reference species (with values ranging from 25% to 76%; Table 1). The genomic compaction of non-coding DNA in genomes A and B can be explained by the shortness of most introns (Figure S4). A specific class of introns with lengths around 25-30 bp (mode at 28 bp) represents 71-72% of the introns. The donor and acceptor sites of these small introns have specific consensus patterns (Figure S4) which are different from other *Porospora* introns. Specifically, these introns exhibit a strongly conserved adenine located 6 bp upstream of the 3’ acceptor site which could represent the intron branch point, as observed for the small introns (20bp) in *B. microti*^31^.

#### Loss of organellar genomes

Recent studies suggest that organellar genomes are lost in most gregarines^10,32^. A precise protocol was set up to identify putative contigs associated with organellar genomes in *P. gigantea*. All the assembled contigs (assigned to *P. gigantea* or not) were searched for regions similar to known organellar genomes. A sensitive protocol based on TBLASTX identified 108 putative regions that were aligned to the NCBI NR library. 102 regions were discarded as bacterial contamination. The 4 contigs corresponding to the remaining 6 regions with at least one significant hit against an eukaryotic sequence were manually curated. Two contigs were assigned to host-derived contaminants whereas the two other long contigs (L=24892 and L=33594) corresponded to *P. gigantea* nuclear genome. Thus, our analyses did not reveal any putative contigs compatible with mitochondrial or apicoplastic genomes.

### Evolutionary histories of *P. cf. gigantea*

#### Genomes A and B diverged several million years ago

We estimated the putative divergence time of A and B genomes by using the divergence between *P. falciparum* and *P. reichenowi* as a calibration point. The synonymous divergence (dS) was calculated for 1003 quartets of orthologous genes. The mean dS value observed between *P. falciparum* and *P. reichenowi* orthologues was 0.0959, similar to that calculated by Neafsey et al^33^ (0.068 substitutions per site) or Reid et al^34^ (0.086-0.11 per site). We assumed that these *Plasmodium* species diverged between 3.3 and 7.7 Mya (TimeTree). The mean dS value observed between the same orthologues in both *P. cf. gigantea* genomes was about 0.4295 substitutions per site. Assuming similar substitution rates in gregarines and *Plasmodium* species, we dated the split between genomes A and B to have occurred between 15.5 Mya and 37.7 Mya. This order of magnitude is similar to the estimation of when the basal splits of the mammal *Plasmodium*^35^ (12.8 Mya) or all *Plasmodium*^36^ (21.0–29.3 Mya) occurred, but is significantly later than the emergence of Nephropidae (lobster family) around 180 Mya^37,38^.

#### Expanded superfamily of crustacean gregarines

To assess the position of *P. cf. gigantea* A and B within Apicomplexa, we constructed a genome-wide phylogeny based on 312 concatenated proteins from the datasets published by Salomaki et al, 2021^13^ and all recently published transcriptomic data from gregarines^10,11,13^ (Figure 4). This phylogeny grouped *P. cf. gigantea* A and B into one clade, placed as a sister group of other crustacean gregarines (*Cephaloidophora communis*, *Heliospora caprellae*), although having shorter branch lengths. In agreement to Salomaki et al (2021)^13^ *Cryptosporidium* species remain at the base of A+G (Apicomplexa + gregarines), using a LG+C60+G+F model in maximum likelihood phylogenomic analyses. However, the bayesian analysis using classical partitioned model LG+G+F is in favor of a A+C topology (Apicomplexans + *Cryptosporidium*) (average standard deviation of split frequencies = 0.020977). More sampling of *Cryptosporidium* relatives is required to address the apicomplexan topology issue.

**Figure 4.**
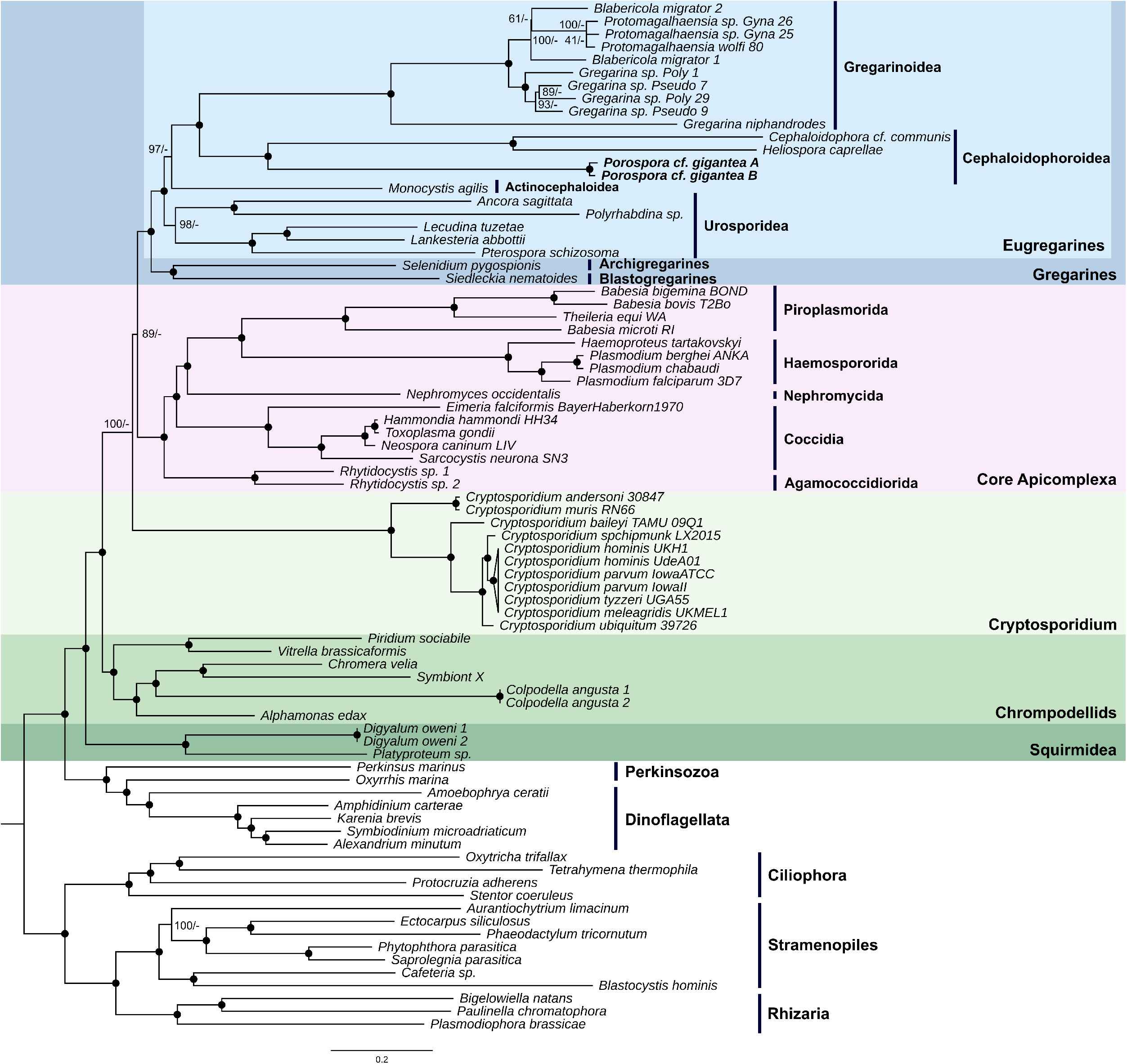
Phylogeny of Apicomplexa. Maximum likelihood phylogeny of apicomplexans as retrieved from a 312 proteins dataset, merged from two previously published datasets^10,11,13^. Final concatenated alignment comprised 93,936 sites from 80 species. Bootstrap support values (n = 1000) followed by MrBayes posterior probabilities are shown on the branches. Black spots indicate 100/1 supports. *Porospora cf. gigantea* A and B sequenced in this study are bolded. See also Figures S6 and S7.

The sequences of 18S small subunit ribosomal DNA, for which the largest taxonomic sampling for gregarines is available in databases, was also used to position *P. cf. gigantea* within the crustacean gregarines. Using a combination of amplifications with specific primers (initially based on Simdyanov et al. (2015)^39^ and Schrével et al. (2016)^40^ then partly redesigned (Figure S5, Table S7)) and *in silico* clustering, we were able to fully reconstruct complete ribosomal loci covering 18S-ITS1-5.8S-ITS2-28S (5977bp) for both A and B genomes. Thirty polymorphic positions were found between A and B, only one within the 18S sequence, and 29 within the 28S sequence (Figure S5). Two phylogenetic studies were performed, one excluding environmental sequences (Figure S6), the other including them (Figure S7). Most environmental sequences are derived from marine sediments from a wide range of habitats but only two sequences are from the North Atlantic where European and American lobsters live.

Congruent with the concatenated phylogeny (Figure 4), both 18S phylogenies assigned *P. cf. gigantea* A and B to their own clade, placed as a sister group to all other crustacean gregarines (*Cephaloidophora*, *Heliospora*, *Thiriotia*, and *Ganymedes* species), as established in Rueckert et al (2011)^41^ (Figures S6 and S7). Five main clades constituting the superfamily Cephaloidophoroidea were retrieved. The four clades previously outlined^41^, redenominated as Ganymedidae, Cephalodophoridae, Thiriotiidae (as proposed by Desportes and Schrével (2013)^6^), and Uradiophoridae, had at their base the clade Porosporidae. Historically defined as the family gathering *Porospora* and *Nematopsis* genera^6^, this clade is constituted of the two sequences of *P. cf. gigantea*. A new putative clade was formed by the five sequences from a Slovenian karst spring published by Mulec and Summers Engel (2019)^42^ (Figure S7), and it is very well supported to be a sister group to four of the crustacean gregarine families, while the family Porosporidae retains its position as a sister group to all these other clades.

### Partially conserved glideosome machinery

We conducted an inventory of the presence or absence of genes encoding proteins involved in the gliding motility based on the molecular description of the so-called glideosome machinery, grouped according to their function as established by Frénal et al (2017)^26^ (Figure 5.A, all orthologues for *P. cf. gigantea* are detailed in Table S8). Genes for these *T. gondii* and *P. falciparum* reference proteins were searched for in both *P. cf. gigantea* genomes and in the genomes of a selection of representative species, as well as the recently published gregarine transcriptomes^10,11,13^.

**Figure 5.**
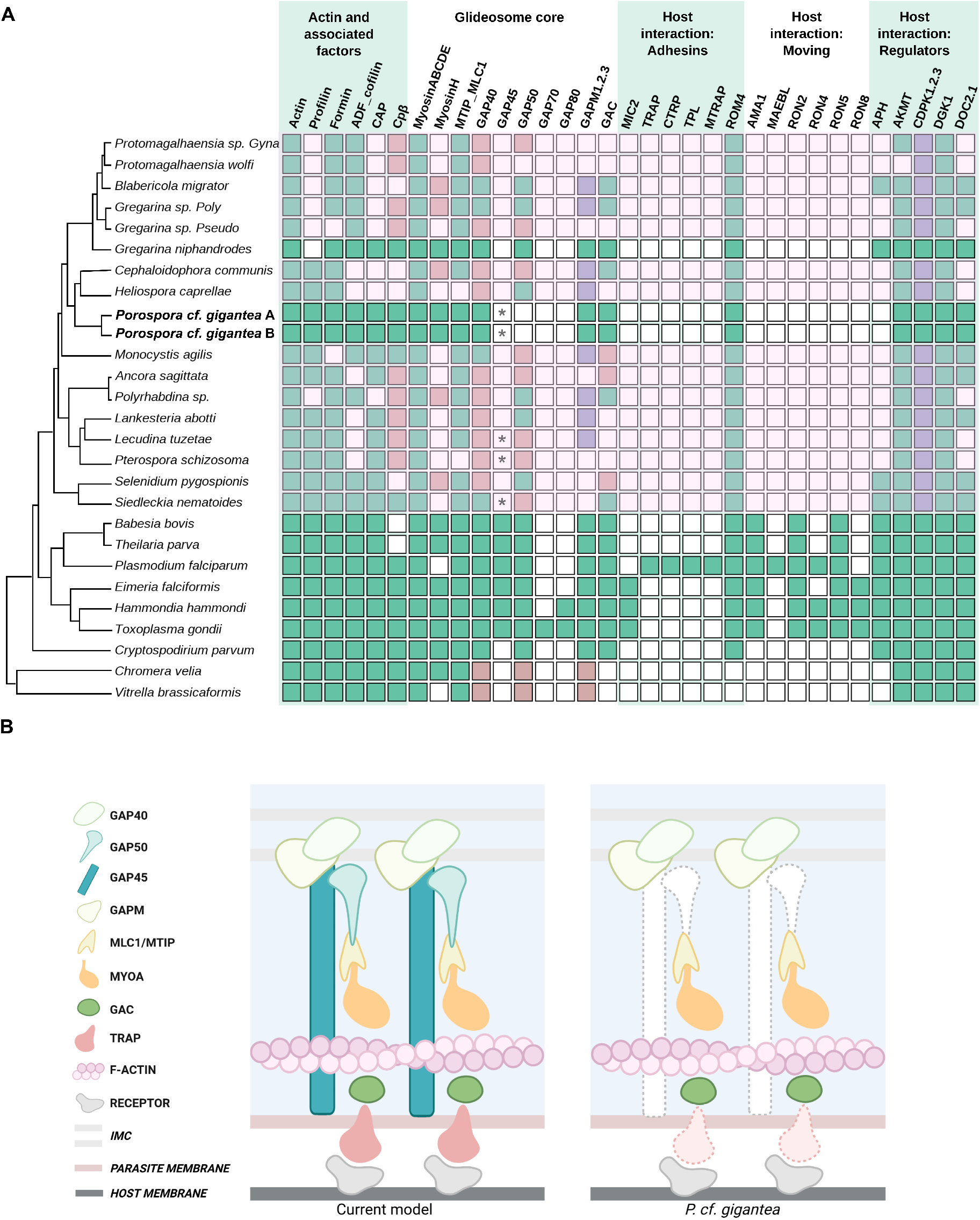
Comparative analysis of glideosome components. **A. Table of presence/absence of genes encoding glideosome proteins, distributed into functional groups.** Glideosome components have been described mainly in *T. gondii* and *P. falciparum*. Protein sequences were searched for in the genomes of both *Porospora* and a selection of representative species as well as in available gregarine transcriptomes. Green indicates the presence, while white indicates the absence of an orthologous protein-encoding sequence. Light red refers to cases where only partial sequences have been retrieved. Violet indicates the presence of at least one protein in multigenic family proteins. * refers to the GAP45 3’ short conserved domain found in some gregarines species. All *P. cf. gigantea* orthologous proteins are detailed in Table S8. **B.** Schematic comparison of the canonical model of the glideosome and the elements found in *P. cf. gigantea* A and B. Missing proteins are shown with dotted lines.

#### Actin and associated factors

Actin in apicomplexans is characterized by a globular monomeric form (G-actin) which polymerizes as needed into short unstable filaments (F-actin)^43^ using various regulators such as profilin^44–46^, ADF cofilin^47^, formin^48–50^, cyclase-associated proteins (CAP)^51^ and F-actin capping protein Cpβ^52^. The inactivation of actin or its associated regulators compromises motility and host cell invasion and egress, although motility may persist in an altered form for a few days, perhaps through alternative mechanisms^26,53–55^. Overall, these proteins are well conserved among Apicomplexa. However, profilin appears to be absent in insect-infecting Gregarinoridea; CAP and Cpβ also seem to be poorly conserved in gregarine transcriptomes but present in both *P. cf. gigantea*.

#### Apicomplexan-specific glideosome proteins

The core glideosome machinery mainly comprises specialized proteins found only in apicomplexans. The single-headed short heavy chain myosin class XIV, named myosin A (MyoA), acts as a motor generating the rearward traction required for gliding motility, invasion and egress, as evidenced by various conditional depletion experiments^56–58^. The glideosome itself is situated between the plasma membrane and the apicomplexan-specific inner membrane complex (IMC). In the IMC, MyoA is associated with a light chain, myosin light chain 1 (MLC1) in *T. gondii* or MyoA tail domain-interacting protein (MTIP) in *P. falciparum*^59^, as well as several glideosome associated proteins (GAP), GAP40, GAP45, GAP50^60–62^, GAP70 and GAP80 as yet only described in *T. gondii*^57^. GAP45 is thought to anchor the glideosome to the plasma membrane by recruiting MyoA as a bridge^62^, whereas GAP40 and GAP50 are predicted to help anchor MyoA to the parasite cytoskeleton^63^. Another set of glideosome-associated proteins with multiple-membrane spans (GAPM) are believed to interact with the alveolin and subpellicular microtubules network, suggesting an indirect interaction with the IMC^26,64^. Finally, the conoid-associated myosin H is necessary for initiating gliding motility in *T. gondii*^65^.

Genes encoding myosins A, B, C, D and E and associated light chains were found in all species. Myosin H is also widely conserved in intracellular apicomplexans. However, among the gregarines Myosin H is only present in a few species. For glideosome associated proteins, only GAP40 was found in all species, although the sequences from gregarine transcripts and chromerids were less well conserved. Surprisingly, given the central role attributed to GAP45 in the glideosome model, no ortholog was found in gregarines except for two poorly conserved sequences in *Lankesteria abotti, Lecudina tuzetae, Cryptosporidium* and chromerids. However, we identified a short conserved 3′ domain (<50aa) in *L. tuzetae, Pterospora schizosoma* and *Siedleckia nematoides*. A similar domain is found in *P. cf. gigantea* A and B. It is however not sufficient to conclude whether it is an orthologous protein. GAP50 seems to be more conserved among apicomplexans, but is absent or only partially conserved in most of the gregarines. As expected, GAP70 and GAP80, only identified so far in *T. gondii*, were not found in other species, except for an orthologue of GAP80 in the coccidia *Hammondia hammondi*. Concerning GAPMs, we found orthologues of at least one of its variants (GAPM 1, 2 or 3) in most species. However, GAPMs seem to be totally absent in at least 7 species of gregarines (*Ancora sagittata, Protomagalhaensia* sp. *Gyna, Protomagalhaensia wolfi, Gregarina* sp. *Pseudo, Pterospora schizosoma, Selenidium pygospionis, Siedleckia nematoides*). Finally, GAC is overall well conserved in apicomplexans but absent from chromerids, supporting its apicomplexan-specific status. However, we were not able to identify GAC in several gregarine transcriptomes (*P*. sp. *Gyna, P. wolfi, G*. sp. *Pseudo, H. capreallae, L. abotti, L. tuzetae, P. schizosoma*) (Figure 4).

#### Adhesins and TRAP-like candidates

The glideosome machinery, anchored in the parasite cytoskeleton, needs to interact with extracellular receptors of the host cell to propel the parasite forward over the host surface. This is made possible by the presence of extracellular adhesins secreted by the micronemes^66,67^ and connected to the glideosome through the glideosome associated connector (GAC) protein^68^. Thrombospondin adhesive protein (TRAP)^69^ is a *Plasmodium* adhesin required for gliding, whose homologue in *T. gondii* is MIC2^70^. At the end of the gliding process, rhomboid protease 4 (ROM4) cleaves the adhesins, disengaging them from receptors and, for intracellular parasites, allowing them to enter the host cell^71–73^. TRAP-like proteins, while highly divergent from one species to another, constitute a family of functionally homologous proteins sharing adhesive domain types, involved in parasite motility and cell penetration^74–76^. TRAP-like or TRAP-related proteins have been detected in various stages of *Plasmodium* (CTRP^77^, MTRAP^78^, TLP^79^) and have also been found *in silico* in *Cryptosporidium* (TRAPCs, CpTSPs^76,80,81^) as well as in several *Babesia* and *Theileria* species^82–85^, in *Neospora caninum*^86^ and in *Eimeria*^87,88^. We first looked for the TRAP proteins which have been implicated in gliding through experimental studies (MIC2, TRAP, TPL, CTRP, MTRAP), as well as the ROM4 protein involved in adhesin cleavage. Unsurprisingly, the currently described TRAP proteins seem to be genus- or even species-specific. On the other hand, we found orthologues for ROM4 in all species, except for chromerids.

The TRAP proteins described to date all have an extracellular region containing one or more TSP1 domains and/or one or more vWA domains^74–76^. They are also characterized by the presence of a single transmembrane domain, a signal peptide, and, in some cases, a juxtaposed rhomboid protease cleavage site, and a short, charged C-terminal cytoplasmic domain with aromatic residues. The presence of a YXXΦ tyrosine sorting signature has also been described^75^ (where X signifies any amino acid, and Φ any hydrophobic amino acid).

To evaluate the presence of TRAP-like proteins in *P. cf. gigantea* genomes, we inventoried all predicted proteins containing at least one TSP1 domain (Table S8), then identified potential candidates with several TRAP-like structural characteristics (Figure 6). We identified a CpTSP2 orthologue within both *P. cf. gigantea* genomes, designated PgTSP2. Like CpTSP2, it is a large protein (~2800 aa) composed of Notch, TSP1, and Sushi domains. PgTSP2 has a localization signal, a transmembrane domain and a short, charged, basic cytoplasmic tail. This protein also has orthologues in *G. niphandrodes*, in chromerids and coccidia.

**Figure 6.**
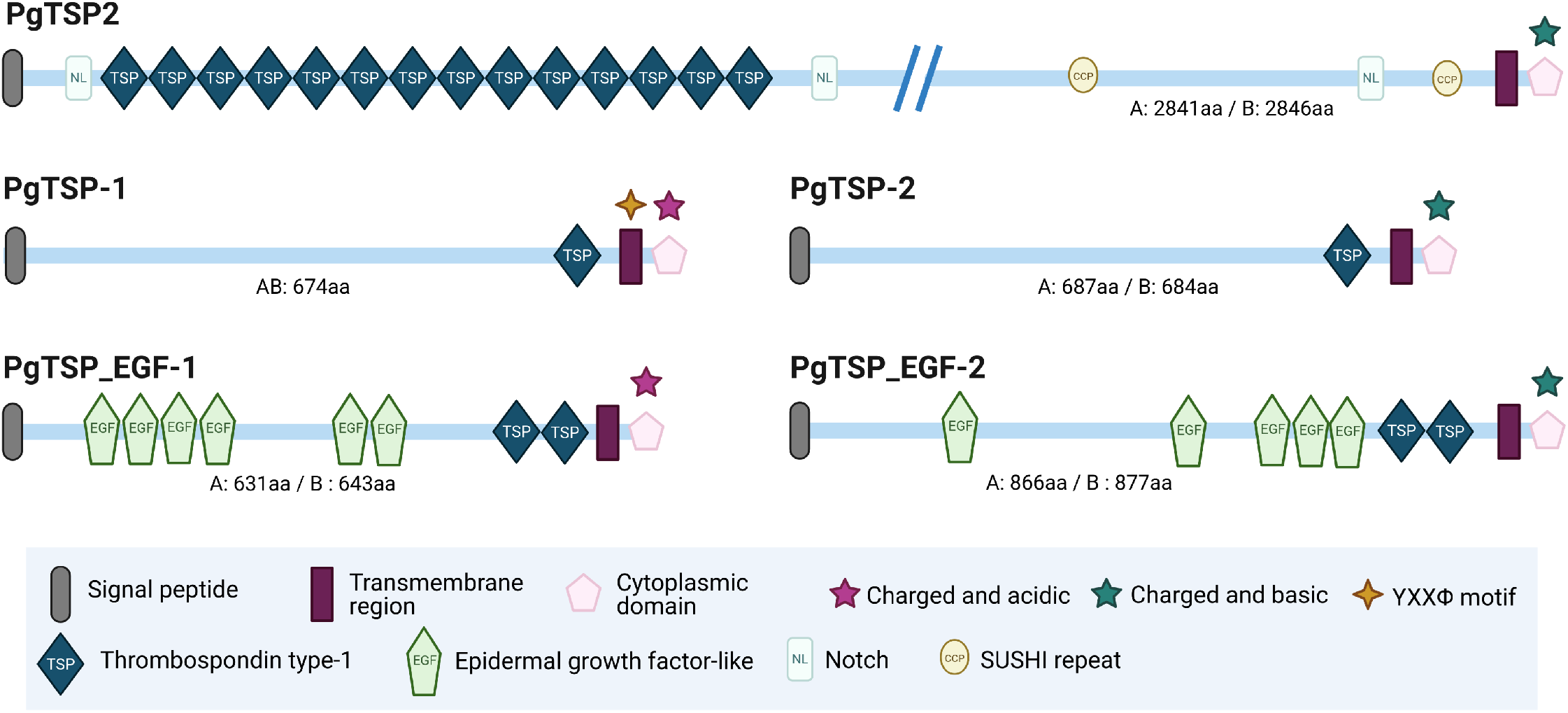
Structures and molecular domains of candidate TRAP-like proteins in *P. cf. gigantea*. **A and B.** See also Table S8.

We demonstrated the presence of genes encoding four other related protein pairs in both A and B genomes, most of which appear to be specific to *P. cf. gigantea*. PgTSP-1 has a TSP1 domain, a signal peptide, a transmembrane domain and a short, charged, acidic cytoplasmic tail. PgTSP-2, very similar in structure to PgTSP-1 also has a TSP1 domain, a signal peptide, a transmembrane domain, and a short, charged but basic cytoplasmic tail. PgTSP_EGF-1 has two TSP1 domains, a signal peptide, a transmembrane domain and a short, charged, acidic cytoplasmic tail, plus several extracellular EGF or EGF-like domains, as also described in *C. parvum* CpTSP7, CpTSP8 and CpTSP9^80^. We identified another protein, PgTSP_EGF-2, very similar in structure.

#### Moving-junction associated proteins

In apicomplexans with intracellular stages such as *T. gondii*, invasion occurs when the tachyzoite initiates a pivotal movement known as reorientation, and a mobile junction settles into the host cell membrane allowing the parasite to enter. Gliding forces are also involved in this process^89^, to which the host cell contributes^90^. A micronemal protein, AMA1, combines with rhoptry neck proteins (RON2, RON4, RON5 and RON8) to firmly secure the parasite to the host cell. In *P. falciparum*, another AMA-like protein, merozoite apical erythrocyte-binding ligand (MAEBL) has an important role in invasion alongside AMA1^91^.

Gregarines remain extracellular during their entire life cycle and *Cryptosporodium* display an intracellular but extra-cytoplasmic stage, so it was not surprising that we did not identify any orthologue of the moving-junction proteins of either these groups. We also searched for predicted proteins implicated in adherence and invasion in *Cryptosporidium*, such as GP15/40, GP900 and mucins, but found no equivalent in either *P. cf. gigantea*^92,93^.

#### Regulatory factors and signaling pathways

Increases in parasite intracellular calcium activate calcium-dependent protein kinases (CDPK) that regulate motility, microneme secretion, invasion and egress^94,95^. Other proteins acting in such signaling pathways include diacylglycerol kinase 1 (DGK1) and acylated pleckstrin homology domain-containing protein (APH), which are involved in microneme secretion regulation^96,97^; the C2 domain-containing protein DOC2.1 which mediates apical microneme exocytosis^98^; and the apical lysine methyltransferase (AKMT), which is involved in gliding motility, invasion and egress in *T. gondii*^99^. We were unable to identify APH in most gregarines and chromerids, and DOC2.1 could not be identified in several transcriptomes. All other regulatory factors appeared to be largely conserved.

## DISCUSSION

### Molecular data support the presence of two species

We report here clear lobster coinfection by two gregarines believed to be distinct that we have named *Porospora cf. gigantea A* and *Porospora cf. gigantea B*. At the molecular level, these two organisms have very similar genomes in terms of size, protein coding capacity, GC content and overall organization with 86% synteny conservation. The delineation of species now requires more precise integrative morpho-molecular approaches, combining extensive imaging (SEM, TEM) and single cell –omics, to find specific traits. Currently, the only molecular tool available for species discrimination in gregarines is the nucleotide sequence of the 18S SSU rDNA. At this molecular marker level, *P. cf. gigantea A* and *P. cf. gigantea B* differ by a single nucleotide, a non-significant divergence for discriminating species.

However, at the genomic level, the genomes show a nucleotide divergence of more than 10% which is incompatible with subspecies or strain definitions. By comparison, applying the same protocol to *P. falciparum* and *P. reichenowi* genomes concluded that the divergence between the two *Plasmodium* species is only 3.2%. Similarly, a divergence of 3-5% between the genomes of *C. parvum* and *Cryptosporidium hominis* has been reported^100^. The large overall genomic divergence between *P. cf. gigantea A* and *P. cf. gigantea B* indicates that they are probably not interfertile, and thus should be considered as different species.

Pending a more integrated morpho-molecular definition of their taxonomy, and better documentation of Cephaloidophoroidea species in general (Figure 4), we propose that *P. cf. gigantea A* and *P. cf. gigantea B* are two distinct organisms infecting *H. gammarus*.

### Two species with compact genomes and a highly specific gene set in common

These two genomes are the first marine gregarine genomes to be sequenced and analyzed and the information generated considerably expands our knowledge of apicomplexan diversity. Both A and B genomes are very small compared to other apicomplexans, with a particularly high gene density. For example, for a similar genome size, *Cryptosporidium* spp. have only about 3900 protein-coding genes compared to the 5300 genes of *P. cf. gigantea*. This result could be partially explained by the absence of certain non-coding sequences in the assemblies, such as centromeres, telomeres and repeated sequences which are difficult to sequence and assemble, notably in *de novo* assembled genomes. However, the compaction is partially due to the comparatively short introns. Small introns with similar consensus sequences have been described in *Babesia microti*^31^.

So far, we have not found any evidence of organellar genomes, whether from the mitochondrion or apicoplast. This needs to be investigated more definitively, especially the mitochondrial aspects. Indeed, the cystic stages from which DNA was collected are unlikely to have many mitochondrial genome copies. To address this issue, it would be more suitable to investigate trophozoite stages via single-cell genomics, for example. According to a recent study, mitochondrial genomes seem to have disappeared from eugregarines^32^. Instead of a distinct mitochondrial genome, the 129 mitochondrial proteins differentially conserved among the gregarine lineages are encoded in the nuclear genome. It would be interesting to identify how many of these nuclear-encoded proteins are conserved within the *P. cf. gigantea* genomes and to reconstruct their specific metabolism. Regarding the apicoplast genome, a recent study stated that it has probably been lost in all eugregarines, while archigregarines may have conserved a highly reduced plastid genome^10^.

BUSCO genome completeness scores of ~70% were found for the two *P. cf. gigantea* genomes, a value not unusual for non-model species^29^, but lower than was found for the *G. niphandrodes* genome (83%) and the 24 other representative species we evaluated (from 76.9% for *Cystoisospora suis* to 100% for *P. falciparum* (Figure S3)). This result also illustrates that the definition of “Apicomplexa core genome” is probably currently highly biased, notably towards *Plasmodium*. Gregarines should be taken into more consideration, as their divergence compared to other apicomplexan models was confirmed by the orthogroup analysis indicating a low percentage of genes conserved between A or B and other studied apicomplexans (<18%).

Even among gregarines the wide diversity is noted as the vast majority of proteins shared by A and B are absent from the *G. niphandrodes* genome. Therefore, studying gregarines will allow a better understanding of the evolutionary history of apicomplexan species, and highlight the astonishing protein diversity brought about by complex differential inheritance from the common ancestor. Through comparative analyses, we will be able to understand how this inheritance has allowed such a wide range of adaptations to parasitism in apicomplexans, which have been able to establish themselves in most Metazoan lineages, vertebrate or invertebrate, marine or terrestrial, in one or more hosts, intracellular or extracellular modes.

### The gregarine glideosome(s)

#### An incomplete but operational machinery

Gliding motility is an essential feature of apicomplexans, and for some intracellular parasites among them, glideosome proteins have been shown to be crucial for host cell invasion and egress^22,23,26,63,74^. However, our sequence analysis of the glideosome components shows that the currently known mechanistic model based on *T. gondii* and *P. falciparum* does not fully account for gliding in all apicomplexans, as anticipated^26,63,67^. Moreover, the conservation of the proteins involved is very variable among the gregarines for which we have omic data. There is little evidence of key molecular components such as canonical adhesins or GAP45, implying that gregarines and *Cryptosporidium* species may have an at least partially alternative machinery dedicated to gliding (Figure 5.B), especially since *P. cf. gigantea* trophozoites are able to glide so rapidly.

#### The model machinery may be partially compensated by alternative proteins

The TRAP adhesin in *T. gondii*, named TgMIC2, has been demonstrated to be an important but non-essential protein to motility^101^. This suggests that TRAP proteins may not be the only proteins involved in host surface adhesion. As we have seen, in the genomes of *P. cf. gigantea* and in other apicomplexans, there are proteins with a structure similar to TRAPs (TRAP-like), that might replace the canonical TRAPs. Understanding the evolution of TRAP requires experimental validation of predicted adhesion proteins in gregarines and *Cryptosporidium* - especially since the presence of these domains in Alveolata does not always correlate with gliding motility^76^. Similarly, the vWA domains, which are found in the canonical TRAPs, appear to be absent from the *Cryptosporidium* genomes. Since gliding is observed in *Cryptosporidium* species, it can be assumed that, if the TRAP-like proteins described in *Cryptosporidium* are indeed involved in gliding, then the vWA domains are not essential for this process. It is also possible that the TSP1 domain proteins represent only one adhesion pathway among others, and that other adhesion domains could perform functions similar to TRAPs, such as the Apple and EGF-like domains in *Cryptosporidium*^75,80^. This is a plausible idea since ROM4, which cleaves adhesins from extracellular receptors of the host cell at the end of the gliding process, is extremely well conserved. GAP45 is thought to maintain the interaction between the IMC and the plasma membrane, and acts as an essential bridge between the two structures^102^. Deleting GAP45 has been proved to prevent glideosome assembly in *P. falciparum*^103^. Perhaps the absence of GAP45 in gregarines and *Cryptosporidium* could be compensated by other GAP-like proteins or it may not even be necessary. Indeed, a looser motor architecture has been proposed, in which actin-myosin motors push in a general backward direction, without necessarily being guided by GAP proteins^63^. Furthermore, while TgMLC1 binding to TgGAP45 is considered a key component of the parasite’s force transduction mechanism, it has recently been shown that loss of TgMLC1 binding to TgGAP45 has little effect on their ability to initiate or maintain movement^104^, questioning again the real role of GAP45 and suggesting our comprehension of the intricacies of the glideosome is still incomplete.

#### Different structures for other forms of gregarine motility

Gregarines have other means of motility, presumably governed by other molecular mechanisms, and the relevance of the glideosome concept to gregarines has been questioned^27,105^. In particular, archigregarines use several modes of movement such as rolling and bending, but not gliding^6,19^. Coelomic and intestinal eugregarines, like crustacean gregarines, have longitudinal, drapery-like surface structures called epicytic folds, the most distinctive feature that differentiates eugregarine trophozoites and gamonts from other apicomplexans. These structures are considered to be involved in eugregarine gliding by increasing the surface area and facilitating actomyosin-based gliding motility, reviewed in Valigurová et al (2013)^27^. Indeed, actin and myosins A, B and F have been localized in epicytic folds in *Gregarina polymorpha*^106,107^. Epicytic folds and mucus, the substance often observed in the trace left by gliding eugregarines^6,27^, are key components to integrate into an alternative model to the current glideosome more representative of eugregarine motility. A particularly interesting study of the crustacean gregarine *Cephaloidophora cf. communis* reported specific attachment apparatus structures^108^. While actin in its polymerized form (F-actin) is observed all along the gregarine, myosin is confined to the cortical region of the cell, in connection with the longitudinal epicytic folds, as described in Valigurová et al. (2013)^27^. This organism also has also a septum, a tubulin-rich filamentous structure that separates the epimerite from the protomerite at the cell apex. Together with microneme-like structures, these features suggest adhesion proteins are produced which could be threaded through the membrane by the numerous pores visible on the epimerite^108^. We were unable to identify alternative movements to gliding in *P. cf. gigantea* (such as peristaltic movement described in other coelomic eugregarines^6,109^). Additional observations are needed to fully document the range of potential motilities in this species, especially since the crustacean-infecting gregarine *C. cf. communis* is capable of jumping or jerking during discontinuous gliding^108^. The different structures invoked, or their absence must be evidenced; indeed, in eugregarines, subpellicular microtubules have never been observed, even though they are supposed to be involved in gliding motility in other apicomplexans^27,108^.

Whatever the molecular mechanisms leading to gliding motility in *P. cf. gigantea*, there are likely to be unique molecular structures, which have evolved consecutive to the specific evolutionary path of gregarines, and which differ from what is currently documented in other apicomplexan lineages.

## ACKNOWLEDGEMENTS

We thank Roscoff Marine Station and service mer & observation for help in collecting hosts; Pauline Konga for the ribosomal genes amplifications; Geraldine Toutirais, Cyril Willig, Marc Gèze and the MNHN Platform (Plateau technique de Microscopie Électronique, Muséum National d’Histoire Naturelle, Paris, France, http://ptme.mnhn.fr/) for the SEM imaging; and the Muséum computational cluster for assemblies/phylogenies.

## AUTHORS’ CONTRIBUTIONS

JB, EDB, LP and IF designed the study. IF, JS and LG performed the biological sampling and IF extracted the nucleic acids. IF, JS and LG performed the photonic microscopy analyses, IF performed the SEM while SLP and GP performed the TEM. JB, EDB and LP did the bioinformatics analyses. AL, IF and LD sequenced and assembled the complete ribosomal loci. JB, IF and EDB performed the glideosome expert annotation. JB performed the 18S phylogenetic analyses. JB, EDB and LP performed the phylogenomic analyses.

JB, EDB, LP and IF wrote the manuscript with contributions from all authors. All authors have read and approved the manuscript.

## MATERIAL & METHODS

### RESOURCE AVAILABILITY

#### Lead contact

Further information and requests for resources should be directed to and will be fulfilled by the lead contact Isabelle Florent.

#### Materials availability

This study did not generate new unique reagents.

#### Data and code availability

DNA and RNA reads and genome assemblies are available in the NCBI database (Bioproject PRJNA734792). Detailled protocols as well as complementary data (phylogenomics datasets, alignments, phylogenetic trees, blasts results and orthogroups) are available on Github at https://github.com/julieboisard/Marine_gregarines_genomes.git/.

### EXPERIMENTAL MODEL AND SUBJECT DETAILS

Several specimens (n = 35) of the lobster species *Homarus gammarus* were collected in the English Channel at Roscoff (Brittany, France) between July 2015 and October 2017 (Table S2), either directly from the wild (Roscoff Bay) or from lobster tank facilities, in which crustaceans are maintained in captivity several weeks to months before their commercialization. According to UICN Red list, *Homarus gammarus* is not an endangered species^110^. The intestinal tract was carefully dissected from each freshly killed host specimen, and transferred to large Petri dishes filled with 0.22-μm filtered, autoclaved sea water, supplemented with the antibiotics penicillin (100 U/mL), streptomycin (100 μg/mL) (Gibco, Life Technologies, USA) and gentamycin (50 μg/mL) (Interchim, Montluçon, France). Trophozoites freely moving in the upper intestine lumen, and cysts loosely attached within the chitinous folds of the hosts’ rectal ampullae (Figure S1), were individually collected using elongated Pasteur pipettes under a classic binocular microscope. For the recording of gliding movement, trophozoites were kept in non-treated sea water. For all other methods, trophozoites, cysts and host tissues were carefully washed several times in 0.22-μm filtered, autoclaved sea water supplemented with the antibiotics indicated above. Trophozoites and cysts were collected for photonic live imaging, scanning electronic microscopy and transmission electronic microscopy, as well as for subsequent omics studies (i.e. DNA and RNA sequencing).

### METHOD DETAILS

#### Electronic microscopy

For the scanning electron microscopy (SEM) studies, isolated trophozoites and cysts, or host intestines and rectal ampullas opened along their longitudinal axis, were washed as indicated above then fixed in 2.5% (v/v) glutaraldehyde in 0.1 M sodium cacodylate (pH 7.2) at 4°C for 6 to 12 hours. After two washing steps in 0.1 M sodium cacodylate (pH 7.2), biological specimens were transferred to microporous specimen capsules (30 μm porosity, 12 mm diameter, 11 mm high, ref #70187-20, Electron Microscopy Science) and dehydrated in a graded series of ethanol in double-distilled water (50, 70, 90, and 100%). Biological specimens in the capsules were critical point-dried in liquid CO_2_ (Emitech K850, Quorum Technologies), then transferred to adhesive carbon-coated holders, and coated with 20 nm of gold (JEOL Fine Coater JFC-1200). Specimens were then examined with a Hitachi SU3500 Premium scanning electron microscope.

For the transmission electron microscopy (TEM) studies, samples were fixed for 2 h in 0.2 M sodium cacodylate buffer with 4% glutaraldehyde, 0.25 M sucrose in 0.2M sodium cacodylate buffer pH 7.4. Cells were then washed three times in sodium cacodylate buffer containing decreasing concentrations of sucrose (0.25 M, 0.12 M, 0 M) for 15 min each time, followed by post-fixation for 1 h at 4°C in 2% osmium tetroxide in 0.1 M sodium cacodylate buffer. After three rinses in 0.2 M sodium cacodylate buffer, samples were dehydrated by successive transfer through an increasing ethanol series (25%, 50%, 70%, 90%, 3 × 100%), then embedded in Spurr’s resin. Sections were cut using a diamond knife on a Leica Ultracut UCT ultramicrotome (Leica, Wetzlar, Germany) and after staining with saturated uranyl acetate for 15 min and Reynolds’ lead citrate for 3 min, were examined on grids with a Jeol 1400 transmission electron microscope (Jeol, Tokyo, Japan).

#### DNA/RNA isolations

Genomic DNA (gDNA) was isolated from 4 biological samples of pooled cysts taken from 3 specimens of the host *H. gammarus:* sample JS-470 from Lobster #7 (~70 cysts), sample JS-482 from Lobster #11 (~50 cysts), samples JS-488 and JS-489 from Lobster #12 (~100 cysts each). Lobster #7 was provided by the Roscoff lobster tank facility while Lobster #11 and Lobster #12 were caught from Roscoff bay. DNA was extracted from the pooled cysts using Macherey Nagel Tissue and Cells isolation kit (ref 740952.50) with yields of 4.1 μg for JS-470, 2 μg for JS-482), 4.5 μg for JS-488 and 6.7 μg (JS-489) of total DNA per sample, as measured by Nanodrop quantification. The protocol was used as recommended by Macherey Nagel, except that the initial lysis step at 56°C was extended beyond the recommended to 1-3 hours with frequent microscopic (binocular) inspection to monitor cyst digestion until completion.

RNA was also isolated from 2 additional biological samples, both composed of pooled cysts taken from the rectal ampulla of their respective hosts: JS-555 (~35 cysts, Lobster #26, Roscoff bay) and JS-575c (~40 cysts, Lobster #34, Roscoff Lobster tank facility). Two distinct protocols were used to isolate total RNA from these two biological samples. For sample JS-555, we used Macherey Nagel basic RNA Isolation kit (ref 740955.10) which yielded ~155 ng of total RNA in 55 μl as assessed by Qbit quantification. For sample JS-575c, we used Macherey Nagel Nucleozol-based RNA Isolation kit (refs 74040.200 and 740406.10) which yielded ~50 ng of total RNA in 55 μl as assessed by Qbit quantification.

#### DNA/RNA sequencing and assembly

The gDNA extracted from the 4 biological samples (JS-470, JS-482, JS-488 and JS-489) was sequenced individually using Illumina NextSeq technology (2 × 151 bp; NextSeq 500 Mid Output Kit v2; Institut du Cerveau et de la Moelle Epinière - CHU Pitié-Salpêtrière - Paris). We obtained 2 × 50 M to 2 × 70 M reads which were checked using FastQC^111^ (version 0.11.5). Reads were cleaned with Trim Galore^112^ (version 0.4.4) which removed remnant Nextera adaptors, clipped 15 bp at 5’-ends and 1 bp at 3’-ends and trimmed low-quality ends (phred score < 30). The assembly was carried out using SPAdes^113^ (version 3.9.1; options: careful mode, automatic k-mers) with the pooled libraries (Figure 2.A).

RNA was extracted from both samples (JS-555 and JS-575c) and treated with RNAse-free DNase. Libraries (Institut du Cerveau et de la Moelle, CHU Pitié Salpétrière, Paris) were prepared following the kit manufacturer’s recommendations (SMART-Seq v4 Ultra Low Input RNA Kit from Takara). Samples were sequenced on a NextSeq 500 Illumina device with MidOutPut cartridge to generate a total of 2 × 87 M reads of 75 bp. The read quality was checked by using FastQC and cleaned by using Trim Galore to remove remnant Nextera adaptors, clipping 15 bp at 5’-ends and 1 bp at 3’-end and trimming low-quality ends (phred score < 30). The sequence reads of both samples were merged into one library which was assembled using Trinity^114,115^.

All genomic contigs longer than 1 kb were analyzed by principal component analysis (PCA) based on their 5-mer composition, which classified them into 6 groups using a hierarchical clustering method (HCA) based on the Ward criterion (Figure 2.B).

For all contigs, the putative protein coding genes were then predicted using Augustus^116^ (version 3.3) and the Apicomplexa gene model for *T. gondii*. All the predicted proteins were thus compared with the NCBI non-redundant protein database using BLAST^117^. The analysis of the taxonomic groups corresponding to the best hits, enabled us to identify five clusters as putative bacterial contaminants whereas the sixth cluster which included 1745 contigs (18.0 Mb), was identified as organisms closely related to Apicomplexa, referred to as the “apicomplexa” cluster (Figure 2.B).

#### Identification of genomes A and B

Preliminary analysis of the “apicomplexa” cluster exhibit two sets of contigs with approximatively 10% of divergence and specific coverage values in the four libraries. The contigs of the “apicomplexa” cluster were split into genomes A and B by using the difference in coverage observed for the four gDNA libraries (Figure 2.C). Each gDNA library (JS-470, JS-482, JS-488 and JS-489) was individually mapped to the contigs using Bowtie2^118^ and the median coverage was calculated for each contig and each library using Samtools^119^ and Bedtools^120^ suites. This coverage information was processed by PCA and a k-means algorithm which classified the contigs into 2 clusters. Then, a linear discriminant model was trained with the coverage information and the result of this first classification before applying it to all the contigs in order to improve the classification. The linear discriminant method (training and classification) was iterated 3 times until convergence. A similar analysis was carried out with 1-kb non-overlapping windows (instead of full-length contigs) to identify putative hybrid contigs. Contigs were thus classified to different genomes depending on the windows, then divided into sub-contigs which were re-assigned to their respective genomes. A detailed protocol with R scripts is available on github (see data and code availability).

The nucleotidic divergence between genome A and genome B was estimated from the alignment of contigs built with Mummer3.0^121^. All alignments of the syntenic regions were parsed to compute the divergence using a home-made script. Assembly metrics were assessed by using QUAST^122^ (version 5.0).

#### Prediction and annotation

All *de novo* assembled transcripts were aligned against the “apicomplexa” cluster contigs with GMAP^123^ within the PASA program^124^. Then, two *ab initio* gene prediction tools, SNAP^125^ (version 2017-11-15) and Augustus were trained using a subset of the PASA transcriptome assemblies. A specific gene model was trained with Augustus, including meta-parameter optimization and prediction of introns (allowing small intron length >10bp) using our “apicomplexan” cluster repeat-masked genome assembly as reference (RepeatMasker^126^, version 4.0.8). Gene predictions were then performed allowing for the prediction of alternative transcripts and noncanonical intron splice sites. An alternative model was also trained with SNAP (default protocol) and used for gene predictions. The Augustus and SNAP outputs showed that some gene predictions were slightly different, so the predictions were parsed with a home-made script to keep as many alternative genes and transcripts as possible for each prediction made. The completeness of the gene prediction was assessed using BUSCO (version 4.0.6).

The predicted proteins were automatically annotated by using i) the best hit of a BLASTP search against VEupathdb (version 2019-20-01), ii) the results of KoFamScam against the KEGG pathway database^127^ (version 2019-05-11) and iii) the signature domains obtained with Interproscan^128^ (version 5.39-77.0).

The ortholog groups were identified with orthoMCL^129^ (default parameters, version 2.0.9) applied to the proteome of a selection of representative organisms available on VEuPathDB (Table S1).

The divergence time of genome A and genome B was estimated from the divergence time of *P. falciparum* and *P. reichenowi* as estimated in TimeTree. Then the coding sequences of the orthologous groups/quartets including a single gene each for genome A, genome B, *P. falciparum* and *P. reichenowi* were aligned using MacSE^130^. For each alignment, the number of synonymous substitutions per site (dS) between genomes A/B and between *P. falciparum/reichenowi* were computed with the maximum likelihood method of Yang and Nielsen (2000)^131^ implemented in PAML4^132^.

The Infernal software^133^ (version 1.3.3) and the Rfam database^134^ (version 14.2) were used together to search for transfer RNAs, spliceosomal RNAs and ribosomal RNAs. The snoReport software^135^ (version 2) was used to search C/D and H/ACA small nucleolar RNAs.

#### Removal of contaminant sequences

##### Host contaminants

All “apicomplexa” cluster contigs were screened against the short reads available from the *Homarus americanus* (PRJNA486050) genome sequencing project, to identify closely-related host contaminants. This dataset was assumed to be free of sequences from apicomplexans, since it was obtained from DNA extracted from non-intestinal tissues (tail, leg or pleiopod appendices). Mapping was carried out with Bowtie2 and the coverages were calculated by using Samtools. The contigs thus identified that were covered over more than 60% of their length by *Homarus* short reads, were considered as host contaminants and were removed.

##### Prokaryotic and fungal contaminants

In parallel, predicted genes in the “apicomplexa” cluster contigs were deeply analyzed for the presence of bacterial and fungal sequences. For each scaffold containing at least one predicted protein, a BLASTP against the NCBI NR database was launched. If the resulting hit had an evalue lower than 1e-30 and more than 30% of the length of the contig was covered by prokaryote/fungi hits, an additional BLASTN against NCBI NR/NT was performed. For the remaining scaffolds without predicted proteins, a direct BLASTN vs NR/NT search was performed. At the end of this procedure, the contigs with prokaryotes/fungi hits covering more than 70% of the length were labeled as contaminants and were removed from the genome assembly.

#### Search for organellar genomes

Organellar genomes were searched using the mitochondrial or apicoplastic genomes available in VEupathDB (version 2019-20-01) as well as with the contigs described in Janouškovec et al. (2019)^10^ as reference sequences. Firstly, a similarity search using a TBLASTX and these sequences as query was applied on all assembled contigs (identified as *P. gigantea* or not). All hits with a bit score above 100 were considered as organellar candidates and were extracted (with 100bp upstream and downstream). Secondly, these candidates were used in a reciprocal TBLASTX search against NCBI NR database to eliminate bacterial contamination. The regions exhibiting at least one hit against an eukaryotic sequence among the nine best hits were manually studied to check if the associated contigs could correspond to organellar genomes.

#### Experimental reconstruction of 18S/28S loci

First, a partial SSU rDNA locus was amplified by using JS-470 gDNA (i.e. genome A only) as template and WL1 and EukP3 primers (Table S7) in a conventional PCR reaction. The amplified bands were cloned and sequenced as previously described^40^. The resulting partial SSU rDNA sequence was further extended in the 3’ direction still using JS-470 gDNA as template and novel primers designed or re-designed based on the molecular data published for *Cephaloidophora cf. communis* and *Heliospora cf. longissima*^39^ (Figure S5A). The resulting sequence (>4 kb) was then used as anchor to reconstruct a complete ribosomal locus with the program iSeGWalker^136^. By clustering reads from JS-470 on this anchor, a 7322-bp theoretical sequence that corresponded to [partial 28S – 18S – ITS1 – 5.8S – ITS2 – partial 28S] including a perfect 1352-bp overlap between the 5’ and 3’ [partial 28S] segments was obtained. From this a complete ribosomal locus [18S – ITS1 – 5.8S – ITS2 –28S] of 5977 bp for genome A was reconstructed, which was validated by PCR amplification, cloning and sequencing (Figure S5B). In a similar clustering approach using all reads for JS-482, JS-488 and JS-489, the complete ribosomal locus for genome B was reconstructed *in silico*, which is the same length but has 30 polymorphisms compared to the genome A locus (Figure S5C). Next, 50% of the complete ribosomal locus for genome B was confirmed by PCR amplification, cloning and sequencing (positions 1187 to 4220, covering partial 18S-ITS1-5,8S-ITS2-partial 28S). This second round of clustering was also used to quantify the respective distributions of genomes A and B present in the latter three biological samples at the full ribosomal locus level. The validated sequence of 18S/28S was manually added to the genome assemblies of genomes A and B, respectively. Schematic representation of rRNA loci was done using BioRender (biorender.com).

#### Phylogeny

##### Phylogenomics of gregarines

The phylogenomic tree was built from a super matrix of 312 orthologues from two datasets published by Salomaki et al (2021)^13^. These two datasets are composed by 246 and 299 orthologues respectively. For all orthologues, corresponding genes have been searched in the proteomes of *P. cf. gigantea* A and B by using BLASTP and candidates were aligned with known orthologues using mafft^137^. Then, orthologous relationships were validated by visual inspection of all the single-protein phylogenetic tree using RaxML^138^ with rapid bootstraps (-f a), -m PROTGAMMAAUTO. Orthologues for *P. cf. gigantea* A and/or B have been recovered for 201 and 256 orthologues in both initial datasets. Both datasets were grouped into a larger dataset composed by 312 non-redundant orthologues. All orthologues were I) filtered with Prequal^139^ to remove non-homologous residues, ii) aligned with mafft, iii) filtered with divvier^140^ to remove alignment errors, iv) trimed with trimAl^141^ and v) merged into the super matrix by using the script *matrix_constructor.py* available with PhyloFisher^142^. The maximum likelihood tree was built with IQ-Tree2 under LG+C60+G+F^143^. The reliability of the phylogenetic tree was tested by the SH-aLRT and ultrafast bootstrap methods (repeated 1 000 times). Bayesian phylogenetic tree was constructed with MrBayes^144^ (version 3.2.3) using a LG+G+F model on a partitioned alignment: prset applyto=(all) aamodelpr=fixed(lg); prset applyto=(all) statefreqpr=fixed(empirical); lset applyto=(all) rates=gamma; unlink shape=(all) pinvar=(all) statefreq=(all); mcmc ngen=500000 samplefreq=1000 printfreq=10000 nchains=4 nruns=2 savebrlens=yes; sump burnin=25000; sumt burnin=25000 contype=allcompat. All trees were visualized and edited using FigTree^145^ (version 1.4.4) and Inkscape (www.inkscape.org).

##### 18S phylogeny of gregarines

The 100-sequence phylogeny was built from the 18S SSU rDNA sequences of the two genotypes of *P. cf. gigantea*, which were aligned with 84 sequences from a diversity of gregarines, either marine or terrestrial, as well as 12 other apicomplexan sequences. Two chromerid sequences were used as the outgroup^146^ but several trees including more than 20 sequences selected from a large diversity of outgroups (from Cryptosporidians, Coccidians, Hematozoans, Colpodellids, Chromerids, Perkinsids, Dinoflagellates, Ciliates, Colponemids, Heterokonts and/or Rhizaria) were built based on Schrével et al (2016)^40^ and conducted to the same conclusions. A total of 1614 sites were found to be conserved after selecting conserved blocks as defined by Gblocks^147^ (version 0.91b) with the following parameters: minimum number of sequences for a conserved position, 51; minimum number of sequences for a flanking position, 51; maximum number of contiguous non-conserved positions, 8; minimum length of a block, 3; allowed gap positions, all. A general time reversible (GTR) substitution model with gamma-distributed rate variation across sites and a proportion of invariant sites was suggested as the best-fit model according to the Bayesian information criterion (BIC) and the Akaike information criterion (AIC) calculated by MEGA X^148^. Maximum likelihood analyses were performed using RAxML (version 8.2.12) with bootstraps estimated from 1,000 replicates. A Bayesian phylogenetic tree was constructed with MrBayes (version 3.2.3) using the following parameters: lset nst = 6 rates = invgamma; mcmc ngen = 10000000, relburnin = yes burninfrac = 0.25, samplefreq =1000, printfreq = 10000, nchains = 4, nruns = 2, savebrlens = yes; sump burnin = 2500000; sumt burnin = 2500000, contype = allcompat.

##### Environmental 18S phylogeny focused on crustacean gregarines

The 189-sequence phylogeny was built from the 18S SSU rDNA sequences from genomes A and B aligned with 14 from crustacean gregarines, and 154 environmental sequences from several projects described in Rueckert et al. (2011)^41^ or gathered from NCBI Genbank. The sequences from the Gregarinoidae clade (n = 19) were used as the outgroup, as this clade has been placed as a sister group to the crustacean gregarine clade in recent literature^10–12^. A total of 1135 sites were found to be conserved after selecting conserved blocks as defined by Gblocks with the following parameters: minimum number of sequences for a conserved position, 95; minimum number of sequences for a flanking position, 95; maximum number of contiguous non-conserved positions, 8; minimum length of a block, 3; allowed gap positions, all. Maximum likelihood and Bayesian analyses were performed following the same protocol and parameters as in the previous 18S phylogeny.

#### Expert annotation for glideosome proteins

A reference apicomplexan glideosome protein dataset was written based on glideosome protein repertoires described in the literature mainly for *T. gondii* and *P. falciparum*^26,63,67^. This reference dataset was used as a seed for parsing the orthogroups established for 25 reference proteomes (Table S1) and the predicted proteomes of the two *P. cf. gigantea* genomes. These reference proteomes were selected by considering the most recent data and associated publications to have the most complete panorama of apicomplexan proteins and key functions/structures documented to date. We also searched for potential orthologues within all recently published proteomes of gregarines using BLASTP (seed: reference proteins in *T. gondii* and *P. falciparum*).

For each orthogroup containing at least one of the reference proteins, the list of proteins was extracted, and the protein sequences were recovered with their respective coding sequences for both *P. cf. gigantea* genomes. BLASTP was performed for extracted proteins against the proteomes of *P. cf. gigantea*, as well as for the candidate proteins from each *P. cf. gigantea* genome against the 25 species reference proteomes. BLASTN was performed against NCBI NR for the coding sequences of the candidate proteins of both *P. cf. gigantea* genomes. The sequences thus collected for each described protein were aligned with mafft. Maximum likelihood molecular phylogeny was deduced from each alignment using RAxML. Analyses were performed using the LG model; bootstraps were estimated from 1,000 replicates. Annotations of the conserved molecular domains were searched for in the automatic annotation and structure analyzed with SMART^149^. For each protein, the results of all the analyses were examined to validate the candidate proteins within the proteomes of the two *P. cf. gigantea* genomes. A table summarizing the presence or absence of glideosome proteins was visualized using R using the tidyverse package^150^. Putative TRAP-like proteins were identified by searching for sequences encoding the TSP1 molecular domain (IPR000884) within the two *P. cf. gigantea* genomes. The predicted structure of each candidate protein was studied, and if necessary partially predicted proteins were re-edited with Genewise^151^. Schematic representation of TRAP-like proteins was done using BioRender (biorender.com).

