## Supplemental Data for "Marine gregarine genomes reveal the breadth of apicomplexan diversity and provide new insights on gliding motility"

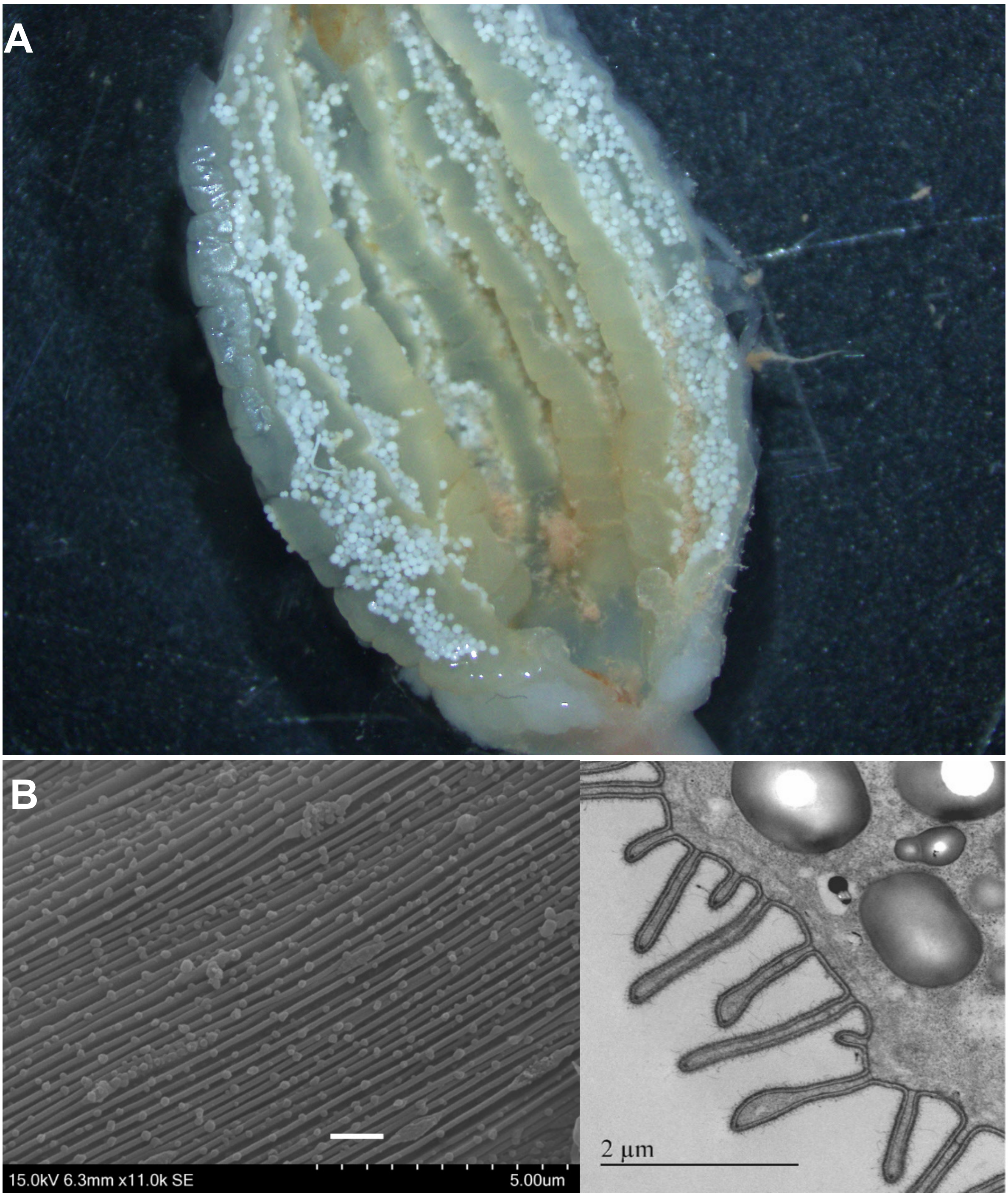

**Figure S1. Additional microscopy figures**, related to Figure 1. A. Photonic image of the rectal ampulla of Lobster#12, longitudinally opened and heavily packed with *Porospora cf. gigantea* cysts in chitinous folds. The length of the rectal ampulla is about 3 cm. B. Morphological evidence for epicytic folds. Zoom on epicytic folds for trophozoite#9, Lobster#12. Scale=1µm. SEM imaging (left); TEM imaging (right).

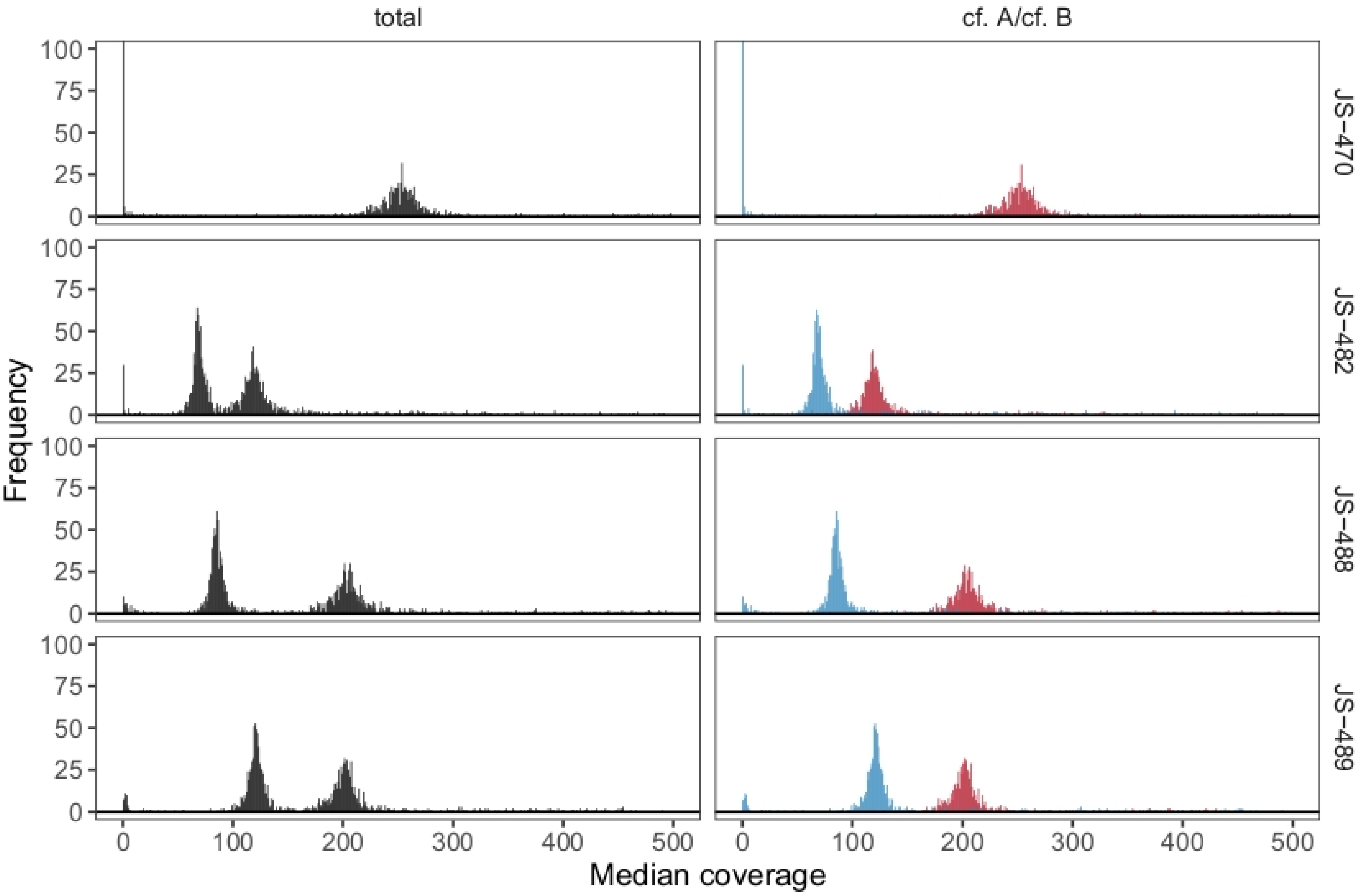

**Figure S2. Distribution of the median coverage in each individual library calculated for each contig from the raw assembly**, related to Figure 2. Total coverage are presented in black (left side). After genomic attribution of each contig, plot is presented again in red for *Porospora cf. gigantea* A and in blue for *Porospora cf. gigantea* B (right side).

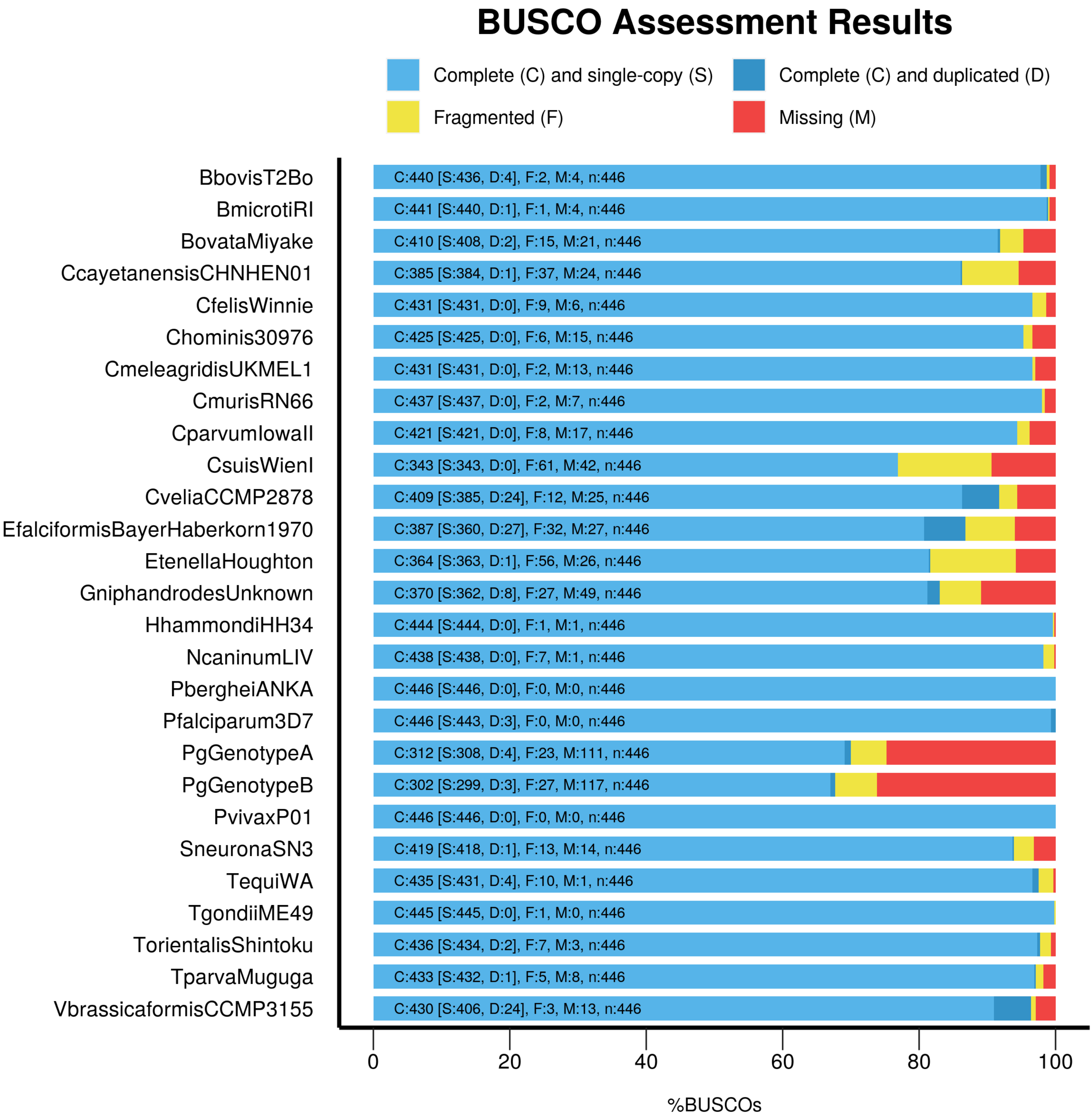

**Figure S3. BUSCOs assessment results for the proteomes of both *P. cf. gigantea* and a selection of 25 reference species** (geneset apicomplexa_odb10), Related to Figure 2.

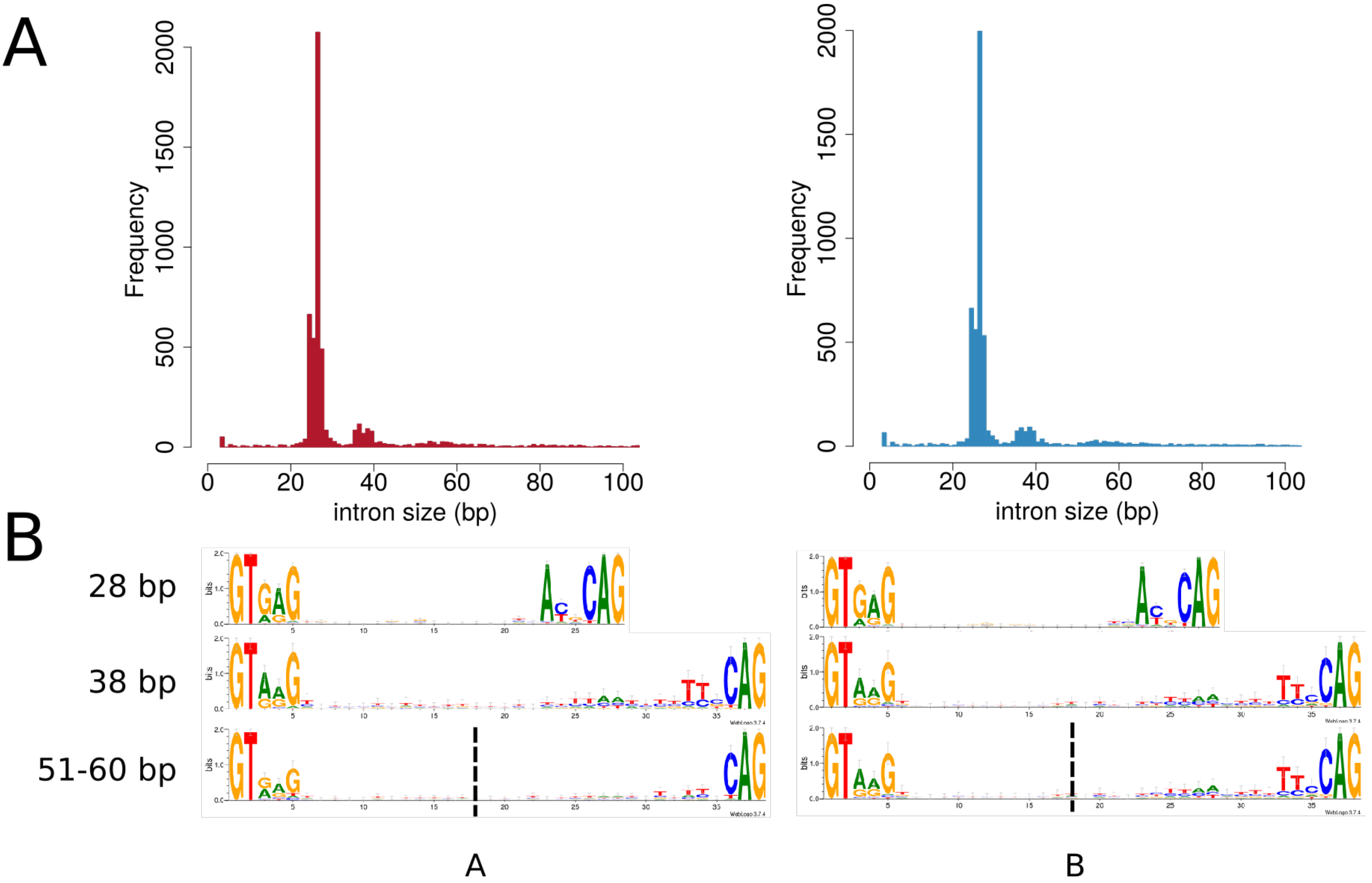

**Figure S4. Introns of both P*. cf. gigantea* genomes**, Related to Table 1 and Figure 2. A. Length distribution B. Consensus for the major class of short introns (28 bp long) and the two other alternative classes. Data for *P. cf. gigantea* A (left side) and B (right side).

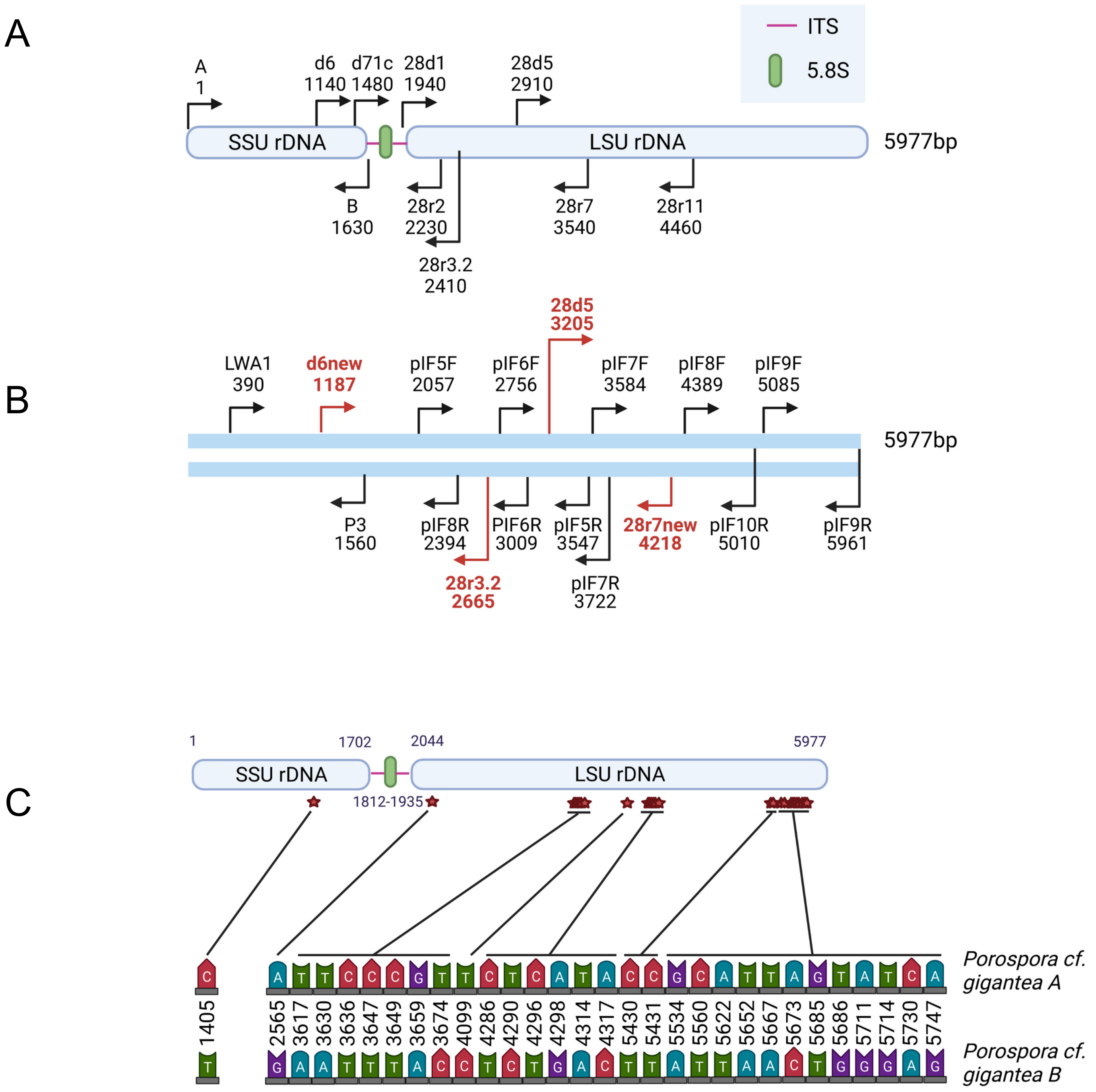

**Figure S5. Complete ribosomal locus reconstruction for *Porospora cf. gigantea*,** related to Figure 2. A. Complete ribosomal locus for *Cephaloidophora cf. communis* and *Heliospora cf. longissima* from Simdyanov et al. (2015)[S1]. B. Complete ribosomal locus for *Porospora cf. gigantea* A using primers based on Simdyanov et al. (2015)[S1] (red) and novel primers (black) to experimentally amplify and sequence the complete 5977bp locus. See also supp. Table 3 for primer sequences. C. Distribution of the 30 polymorphic positions between A and B complete ribosomal loci.

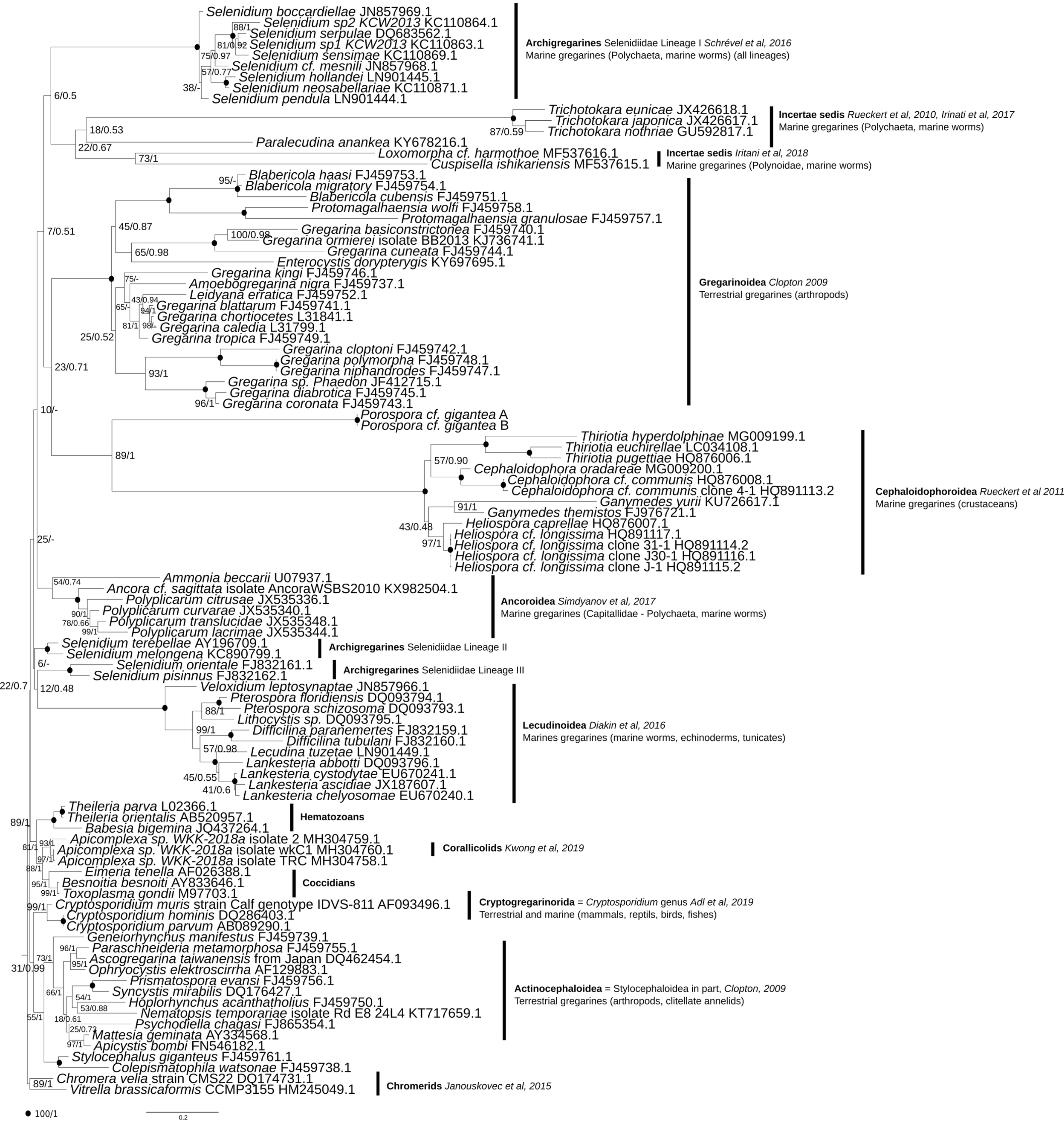

**Figure S6. Gregarines/apicomplexan phylogeny**, Related to Figure 4. Phylogenetic tree built using 100 18S rDNA sequences 1614 sites in order to situate *P. cf. gigantea* A and B among other known gregarines and apicomplexan clades. Chromerid sequences were used as outgroup, as they are considered as the sister group of all other apicomplexans[S2]. Evolutionary history was inferred by maximum likelihood and bayesian inference using a GTR+G+I model. Topologies were identical according to both methods. Black spots indicate 100/1 supports. Supports <70/0.7 are not shown. Families and associated literature are indicated.

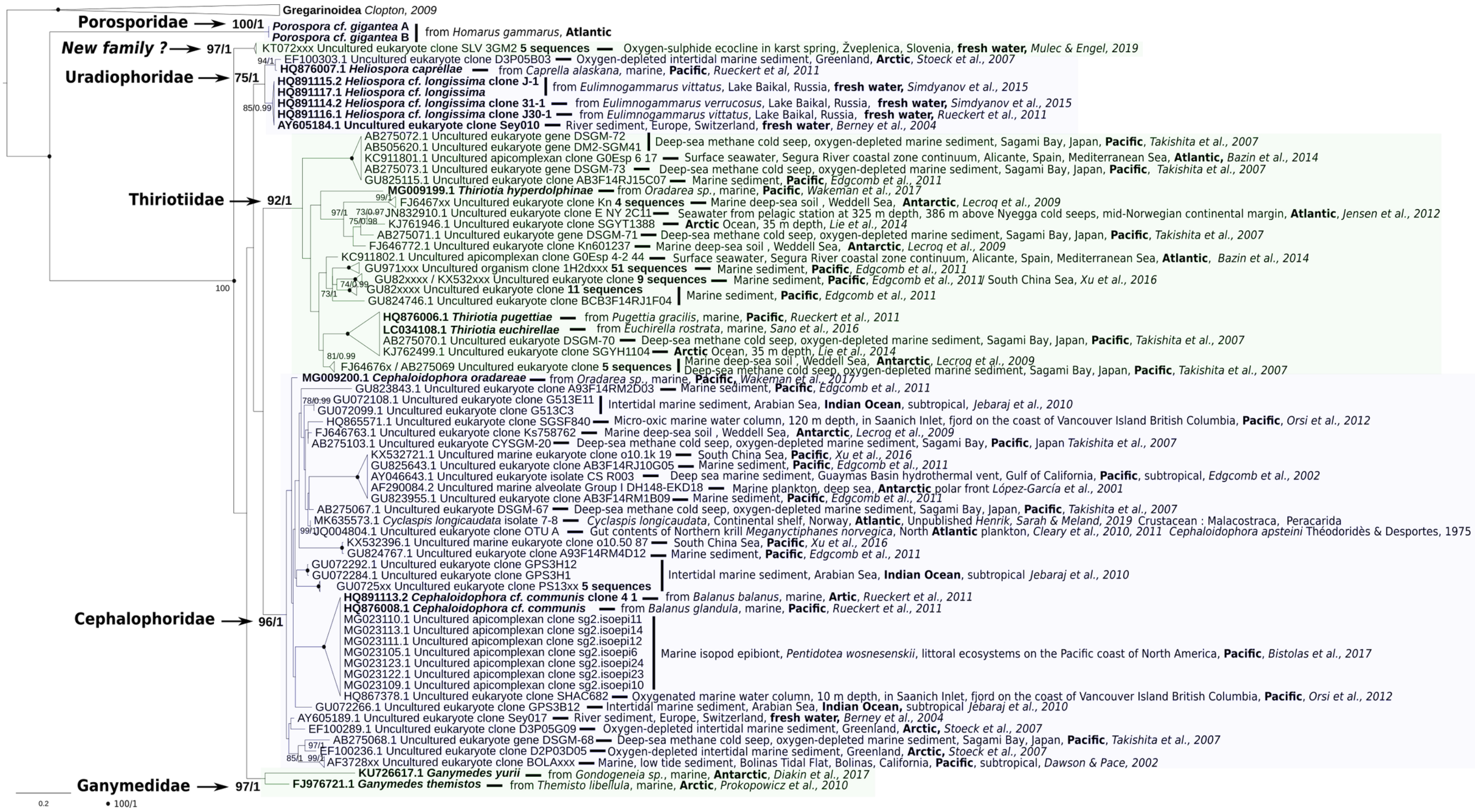

**Figure S7. Environmental phylogeny**, related to Figure 4. Phylogenetic tree built using 189 18S rDNA sequences for 1135 sites in order to situate two *P. cf. gigantea* A and B among other crustacean gregarines and environmental sequences. Considering that Gregarinoidea sequences were placed as sister group of other crustaceans' gregarines in the gregarines/apicomplexan phylogeny, as well as in recent literature[S3,S4,S5], they were used as outgroup. Evolutionary history was inferred by maximum likelihood and bayesian inference using a GTR+G+I model. Topologies are identical according to both methods. Black spots indicate 100/1 supports. Supports <70/0.7 are not shown. Geographical provenance of all environmental sequences are indicated and their localization is highlighted in bold.

| Species | Strain | Gene count (a) | Contigs (a) | Total length (Mb)(b) | GC (%) (b) | Publication (a) |
|---|---|---|---|---|---|---|
| Cryptosporidium hominis | 30976 | 3994 | 53 | 9.059 | 30.13 | Guo et al, 2016[S6] |
| Cryptosporidium muris | RN66 | 3981 | 75 | 9.242 | 28.47 | x |
| Cryptosporidium meleagridis | UKMEL1 | 3806 | 57 | 8.973 | 30.97 | Ifeonu et al, 2016[S7] |
| **Cryptosporidium parvum (1)** | **IowaII** | **4020** | **8** | **9.102** | **30.22** | **Abramhasen et al, 2004**[S8] |
| **Chromera velia (1)** | **CCMP2878** | **30806** | **5953** | **193.884** | **49.11** | **Woo et al, 2015**[S2] |
| **Vitrella brassicaformis (1)** | **CCMP3155** | **23503** | **1064** | **72.700** | **58.09** | **Woo et al, 2015**[S2] |
| **Gregarina niphandrodes (1)** | **Unknown** | **6606** | **468** | **14.008** | **53.78** | **x** |
| Cyclospora cayetanensis | CHN_HEN01 | 7592 | 2297 | 44.034 | 51.84 | Liu et al, 2016[S9] |
| Cystoisospora suis | WienI | 11767 | 7880 | 81.642 | 49.32 | Palmieri et al, 2017[S10] |
| Eimeria falciformis | BayerHaberkorn1970 | 6037 | 753 | 43.672 | 49.86 | Heitlinger et al, 2014[S11] |
| Eimeria tenella | Houghton | 8634 | 4664 | 51.859 | 51.33 | Reid et al, 2014[S12] |
| Hammondia hammondi | HH34 | 8177 | 3676 | 64.338 | 52.83 | Walzer et al, 2013[S13] |
| Neospora caninum | LIV | 7266 | 66 | 59.103 | 54.82 | Reid et al, 2012[S14] |
| Sarcocystis neurona | SN3 | 7089 | 873 | 124.411 | 51.41 | Blazejewski et al, 2015[S15] |
| **Toxoplasma gondii (1)** | **ME49** | **8920** | **2075** | **65.590** | **52.30** | **Lorenzi et al, 2016**[S16] |
| Babesia bovis | T2Bo | 3781 | 14 | 8.179 | 41.59 | Brayton et al, 2007[S17] |
| Babesia microti | RI | 3685 | 6 | 6.434 | 36.17 | Cornillot et al, 2012[S18] |
| Babesia ovata | Miyake | 5108 | 91 | 14.453 | 49.27 | Yamagishi et al, 2017[S19] |
| Theileria equi | WA | 5397 | 12 | 11.674 | 39.48 | Kappmeyer et al, 2012[S20] |
| Theileria orientalis | Shintoku | 4058 | 6 | 9.010 | 41.55 | Hayashida et al, 2012[S21] |
| Theileria parva | Muguga | 4167 | 10 | 8.353 | 34.04 | Gardner et al, 2005[S22] |
| Cytauxzoon felis | Winnie | 4389 | 357 | 9.108 | 31.81 | Tarigo et al, 2013[S23] |
| Plasmodium berghei | ANKA | 5245 | 21 | 18.780 | 22,04 | Otto et al, 2014[S24] |
| **Plasmodium falciparum (1, 2)** | **3D7** | **5712** | **16** | **23.332** | **19.34** | **Gardner et al, 2002**[S25] |
| Plasmodium vivax | P01 | 6830 | 242 | 29.052 | 39.78 | Auburn et al, 2016[S26] |
| Plasmodium reichenowi (2) | G01 | 5909 | 48 | 24.471 | 24.47 | Otto et al, 2014[S24] |

(1) subset of 6 species (4 apicomplexan + 2 chromerids) used in some comparative analyses, including search for orthogroups and genomic metrics
(2) species used to date the divergence of *P. cf. gigantea* A and B
(a) data from VEupathDB release 41[S27]
(b) data obtained with QUAST[S28]

**Table S1. Metrics of 25 apicomplexan and chromerids genomes, considered representative for comparative analyses**. Related to Table 1 and Figure 3.

| Lobster Specimen | Sampling date | Host from Tanks/Bay | Host sex | Host Lenght (cm) | Host Weight (g) | Cysts load in host rectal ampulla | Trophozoites Load in host gut lumen |
|---|---|---|---|---|---|---|---|
| #1 | 24/05/2016 | Tanks | male | 25 | 355 | < 10 | none |
| #2 | 24/05/2016 | Tanks | male | 29 | 645 | 10-100 | none |
| #3 | 24/05/2016 | Tanks | female | 26 | 450 | 10-100 | none |
| #4 | 25/05/2016 | Tanks | female | 29 | 620 | < 10 | none |
| #5 | 25/05/2016 | Tanks | male | 25 | 420 | 10-100 | none |
| #6 | 26/05/2016 | Tanks | male | 29 | 745 | < 10 | < 10 |
| #7 | 26/05/2016 | Tanks | male | 25 | 375 | 10-100 | < 10 |
| #8 | 27/05/2016 | Tanks | female | 27 | 445 | 10-100 | none |
| #9 | 27/05/2016 | Tanks | male | 26 | 490 | 10-100 | none |
| #10 | 30/05/2016 | Tanks | male | 26 | 470 | none | none |
| #11 | 30/05/2016 | Bay | female | 25 | 420 | 10-100 | none |
| #12 | 31/05/2016 | Bay | male | 24 | 465 | 100-1000 | >10 |
| #13 | 31/05/2016 | Bay | female | 24 | 435 | 100-1000 | none |
| #14 | 18/10/2016 | Tanks | male | 27 | 485 | ~200 | < 10 |
| #15 | 19/10/2016 | Tanks | male | 26 | 685 | none | none |
| #16 | 19/10/2016 | Tanks | female | 27 | 535 | none | none |
| #17 | 20/10/2016 | Bay | male | 23 | 455 | 100-300 | < 10 |
| #18 | 20/10/2016 | Bay | male | 25 | 450 | 10-100 | < 10 |
| #19 | 24/10/2016 | Bay | female | 25 | 510 | 10-100 | none |
| #20 | 24/10/2016 | Tanks | male | 23 | 405 | 10-100 | >10 |
| #21 | 25/10/2016 | Tanks | male | 27 | 550 | none | none |
| #22 | 26/10/2016 | Tanks | female | 30 | 895 | 10-100 | >10 |
| #23 | 26/10/2016 | Tanks | male | 29 | 510 | 10-100 | none |
| #24 | 03/10/2017 | Tanks | female | 27 | 505 | 10-100 | >10 |
| #25 | 03/10/2017 | Tanks | female | 28 | 580 | 10-100 | >10 |
| #26 | 04/10/2017 | Bay | male | 34 | 815 | 10-100 | none |
| #27 | 05/10/2017 | Bay | male | 26 | 515 | 100-500 | >200 |
| #28 | 06/10/2017 | Tanks | female | 30 | 635 | < 10 | none |
| #29 | 06/10/2017 | Tanks | male | 26 | 560 | 10-100 | none |
| #30 | 09/10/2017 | Tanks | male | 27 | 655 | none | none |
| #31 | 09/10/2017 | Tanks | male | 28 | 710 | none | none |
| #32 | 09/10/2017 | Tanks | female | 26 | 450 | 10-100 | none |
| #33 | 11/10/2017 | Tanks | male | 26 | 470 | 10-100 | >10 |
| #34 | 12/10/2017 | Tanks | male | 27 | 510 | 10-100 | none |
| #35 | 17/07/2015 | Bay | | | | 100-300 | none |

**Table S2. Sampling of the lobster specimen.** Related to Figure 1.

| Trophozoite specimen | Host specimen (origin) | Length (µm) | Width <u>+</u> SD (µm) (n=number of measures) |
|---|---|---|---|
| #1 | H0 (Bay) | 1796 | 32.8±4.5 (n=13) |
| #2 | H0 (Bay) | >983 | 34.2±3.9 (n=14) |
| #3 | H12 (Bay) | none | 45.2±3.6 (n=3) |
| #4 | H6 (Tank) | 1424 | 51.5±8.4 (n=25) |
| #5 | H12 (Bay) | none | 66 to 23µm |
| #6 | H12 (Bay) | none | none |
| #7 | H12 (Bay) | >1043 | 71.5±10.1 |
| #8 | H12 (Bay) | 1858 | 43.3±7.7 (n=13) |
| #9 | H12 (Bay) | 2585 | 55.5±4.5 (n=6) |
| #10 | H12 (Bay) | none | 41 |
| #11 | H20 (Tank) | 2000 | 36.2±3.1 (n=6) |
| #12 | H20 (Tank) | 2177 | 37.6±7.3 (n=6) |
| #13 | H20 (Tank) | 2222 | 30.6±1.9 (n=6) |
| #14 | H20 (Tank) | >1062 | 51.0±6.6 (n=6) |
| #15 | H20 (Tank) | >681 | 31.5±1.9 (n=6) |
| **Mean value** | | | **41.8+10.4 (n=104)** |

(a) mean values for 15 trophozoites from indicated hosts specimen
(b) The sign > corresponds to truncated trophozoites that could not be measured in full.

**Table S3. Length and width of trophozoites**, related to Figure 1. All values are based on SEM images.

| Cyst specimen | Host specimen (origin) | Diameter (µm) (a) |
|---|---|---|
| #1 | H#12 (Bay) | 118.7±4.5 (n=8) * |
| #2 | H#6 (Tank) | 168.4±9.1 (n=3) * |
| #3 | H#12 (Bay) | 135.3±1.6 (n=4) |
| #4 | H#12 (Bay) | 168.6±2.5 (n=4) |
| #5 | H#12 (Bay) | 157.1±5.4 (n=4) |
| #6 | H#12 (Bay) | 122.7±2.8 (n=4) |
| #7 | H#12 (Bay) | 162.0±4.0 (n=4) |
| #8 | H#12 (Bay) | 120.6±3.6 (n=4) |
| #9 | H#4 (Tank) | 137.0±1.7 (n=4) |
| #10 | H#4 (Tank) | 108.4±10.6 (n=4) |
| #11 | H#4 (Tank) | 109.8±3.9 (n=4) * |
| #12 | H#4 (Tank) | 168.6x128.4 (oval) (n=2) |
| #13 | H#4 (Tank) | 220.6±7.0 (n=4) |
| #14 | H#4 (Tank) | 252.2±3.7 (n=4) |
| #15 | H#4 (Tank) | 240.9±6.5 (n=4) |
| #16 | H#4 (Tank) | 211.0±10.9 (n=4) |
| #17 | H#4 (Tank) | 141.9±2.0 (n=4) |
| #18 | H#4 (Tank) | 118.1±1.4 (n=4) |
| #19 | H#4 (Tank) | 104.6±3.2 (n=4) |
| #20 | H#4 (Tank) | 108.3±3.1 (n=4) |
| #21 | H#4 (Tank) | 121.9±4.1 (n=4) |
| #22 | H#4 (Tank) | 129.7±6.1 (n=4) |
| #23 | H#4 (Tank) | 124.7±3.6 (n=4) |
| #24 | H#4 (Tank) | 230±9.9 (n=4) |
| #25 | H#4 (Tank) | 220.6±7.0 (n=4) |
| **Mean** | | **151.1±45.3 (n=97)** |

(a) Mean values measured for 25 cysts. One diameter ± standard deviation (for spherical cysts) or two measures (for oval cyst #12) are given. n, number of measures.
* cysts that were further investigated for gymnospores and zoites measures (see Supplementary Table 6).

**Table S4. Diameters of cysts**, related to Figure 1. All values are based on SEM images.

| Origin of gymnospores and zoites | Host specimen (origin) | Gymnospore Diameter (µm)(a) | Zoite length (µm) (a) | Zoite width (µm) (a) |
|---|---|---|---|---|
| Cyst#1 | H#12 (Bay) | 4.97±0.37 (n=60) | 1.17±0.07 (n=7) | 0.565±0.218 (n=50) |
| Cyst#2 | H#6 (Tanks) | 5.79±0.62 (n=97) | 1.04±0.11 (n=45) | 0.616±0.033 (n=11) |
| Cyst#11 | H#4 (Tanks) | 6.04±0.72 (n=56) | 1.09±0.07 (n=16) | 0.674±0.043 (n=35) |
| JS-463b_0003 | H#4 (Tanks) | | | 0.660±0.045 (n=10) |
| JS-463b_0016 (a) | H#4 (Tanks) | 5.30±0.05 (n=4) | | 0.643±0.042 (n=10) |
| JS-463b_0016 (b) | H#4 (Tanks) | 5.08±0.16 (n=4) | | |
| JS-463b_0016 (c) | H#4 (Tanks) | 5.92±0.18 (n=4) | 1.19±0.09 (n=3) | 0.673±0.086 (n=3) |
| JS-463b_0020 (a) | H#4 (Tanks) | 5.26±0.18 (n=4) | | 0.662±0.035 (n=10) |
| JS-463b_0020 (b) | H#4 (Tanks) | 5.33±0.04 (n=4) | | |
| JS-463b_0020 (c) | H#4 (Tanks) | 6.21±0.22 (n=4) | | |
| JS-463b_0020 (d) | H#4 (Tanks) | 6.63±0.30 (n=4) | | |
| JS-463b_0020 (e) | H#4 (Tanks) | 6.24±0.18 (n=4) | | |
| JS-463b_0020 (f) | H#4 (Tanks) | 5.64±0.11 (n=4) | | |
| JS-463b_0027 (a) | H#4 (Tanks) | 5.66±0.19 (n=4) | | 0.646±0.050 (n=10) |
| JS-463b_0027 (b) | H#4 (Tanks) | 4.69±0.10 (n=4) | | 0.656±0.029 (n=10) |
| JS-463b_0027 (c) | H#4 (Tanks) | 5.70±0.10 (n=4) | | 0.643±0.039 (n=10) |
| JS-463b_0028 | H#4 (Tanks) | | 1.09±0.09 (n=10) | |
| JS-463b_0030 | H#4 (Tanks) | 5.80±0.08 (n=4) | | 0.684±0.085 (n=10) |
| JS-463b_0036 | H#4 (Tanks) | | 1.16±0.07 (n=13) | 0.613±0.037 (n=10) |
| **Mean** | | **5.63±0.69 (n=265)** | **1.04±0.16 (n=105)** | **0.630±0129 (n=176)** |

(a) n, number of measures.

**Table S5. Diameters of gymnospores and zoites**, related to Figure 1. Diameters were measured for hundreds of gymnospores within cysts (3 first lines) or released from cysts (remaining lines). Whenever possible, length and apical width ± standard deviation of their constitutive zoites were also measured. All values are based on SEM images.

| Video record | Length of recording (s) | Trophozoites number | Length (µm) | Speed (µm/s) |
|---|---|---|---|---|
| G5310002 | 40 | T10<br>T11<br>T12<br>T13<br>T14<br>T15 | ~2190<br>~1876<br>~2113<br>~2113<br>~1801<br>~1807 | 51.6<br>48.9<br>50.3<br>51.8<br>49.8<br>51.4 |
| G5310003 | 34 | T3<br>T4<br>T5 | ~3100<br>~4500<br>~3900 | 87-89<br>100-109<br>108-115 |
| G5310004 | 60 and 48 | T1<br>T2 | ~3000<br>~3600 | 56-63<br>80-81 |
| G5310018 | 20 | T6<br>T7<br>T8<br>T9 | ~2540<br>~4600<br>~4100<br>~3595 | 76<br>97-103<br>104<br>91-94 |

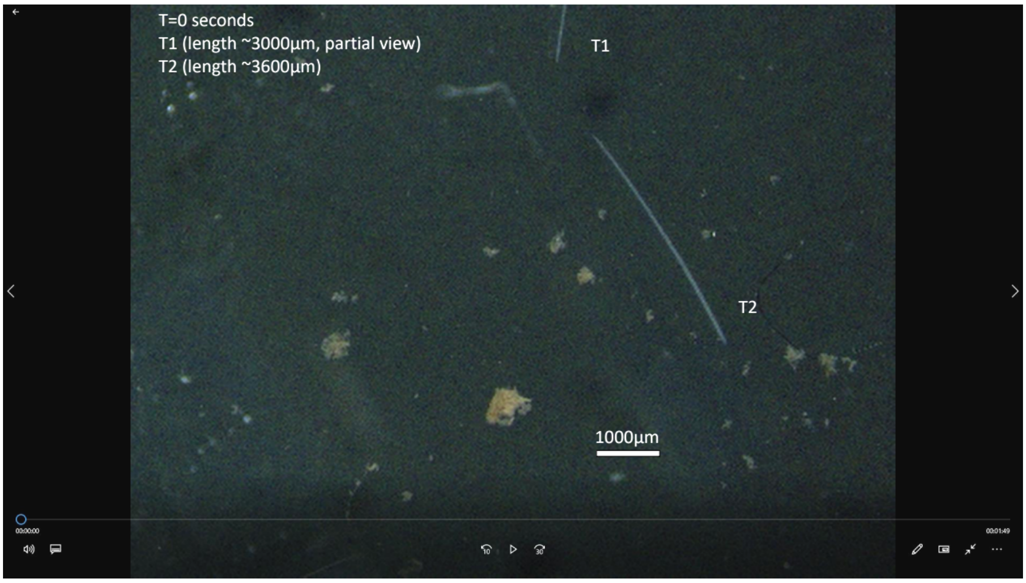

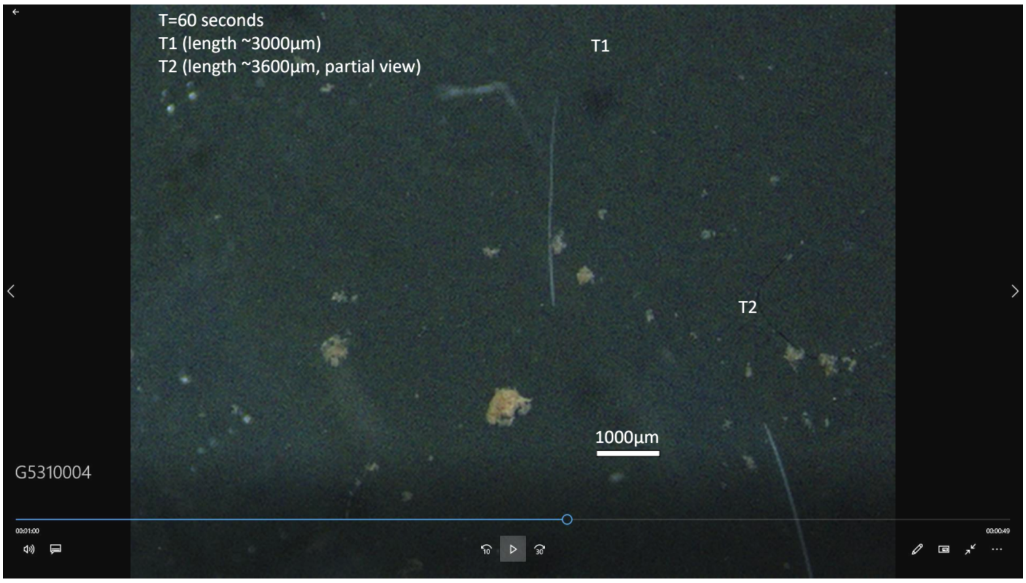

**Table S6. Gliding recordings**, related to Figure 1. All recordings are from trophozoites collected from Lobsters#12 and #13 on 31/05/2016. Due to the lack of scale on these videos, we used the mean width of trophozoites, as determined by using SEM images (41.8±10.4µm, see Table S4), to calibrate the other measures. Two screenshots (t=0 and t=60') from the supplemental film are reproduced.

| Primer name | Primer sequence | orientation | reference |
|---|---|---|---|
| LWA1 | 5'- GGAAGGCAGCAGGCGCGC - 3' | forward | Schrevel et al., 2016[S29] |
| EukP3 | 5'- GACGGGCGGTGTGTAC - 3' | reverse | Lara et al., 2007[S30] |
| 28d5 | 5'- CCGCTAAGGAGTGTGTAACAAC - 3' | forward | Simdyanov et al., 2015[S1] |
| 28r3.2 | 5'- ACTCCTYRGTCCGTGTTTCA - 3' | reverse | Simdyanov et al., 2015[S1] |
| 28r2 | 5'- TACTTGTYBRCTATCG - 3' | reverse | Simdyanov et al., 2015[S1] |
| d6new | 5'- GGTGGTGCATGGCCAAACTT - 3' | forward | Modified from Simdyanov et al., 2015[S1] |
| 28d5short | 5'- GCTAAGGAGTGTGTAACAAC - 3' | forward | Modified from Simdyanov et al., 2015[S1] |
| 28r7new | 5'- TAATTTGCCGACTTCCCTCA - 3' | reverse | Modified from Simdyanov et al., 2015[S1] |
| PIF5F | 5'- ACATTCCTTGGGTTACCC - 3' | forward | This study |
| PIF6F | 5'- TAACGACCCGAAAATCGG - 3' | forward | This study |
| PIF7F | 5'- CATGCTAACACAAGGGGG - 3' | forward | This study |
| PIF8F | 5'- CCGACAGTTTAACTAAAACC - 3' | forward | This study |
| PIF9F | 5'- GAGATCATATCGACGCGG- 3' | forward | This study |
| PIF5R | 5'- CATCAGTGCGACGATACC - 3' | reverse | This study |
| PIF6R | 5'- GTTTGAGAATCAGTCGAGG - 3' | reverse | This study |
| PIF7R | 5'- CTTTCGACTTCCGACAGC - 3' | reverse | This study |
| PIF8R | 5'- TTGTTTGCTATCGGTATAGG - 3' | reverse | This study |
| PIF9R | 5'- AAATCTCAAGAGAGATGGAG- 3' | reverse | This study |
| PIF10R | 5'- GCTAAGGATCGATAGGCC - 3' | reverse | This study |

**Table S7. List of primers used for ribosomal locus amplification and sequencing by Sanger technology**. Related to Figure S5.

| Protein name | *P. cf. gigantea* A | *P. cf. gigantea* B |
|---|---|---|
| Actin | KAH0480873.1<br>KAH0486428.1<br>KAH0477721.1<br>KAH0476702-3.1 | KAH0488956.1<br>KAH0472333.1<br>KAH0482523.1 |
| Profilin | KAH0486108.1 | KAH0488334.1 |
| Formin | KAH0487606.1<br>KAH0473072-3.1 | KAH0488107-8.1<br>KAH0473233-4.1 |
| ADF_cofilin | KAH0487744.1 | KAH0488925.1 |
| CAP | KAH0483375-6.1 | KAH0475361-2.1 |
| Cpβ F-actin capping protein β-subunit | KAH0474592:4.1 | KAH0471516:8.1 |
| MyosinACDE ClassXIV | KAH0483751.1<br>KAH0483531.1<br>KAH0484363.1<br>KAH0481456.1<br>KAH0475489.1 | KAH0487710:12.1<br>KAH0480515.1<br>KAH0484061.1<br>KAH0483307:11.1 |
| MyosinH ClassXIV | KAH0473906:8.1 | KAH0486610:12.1 |
| MTIP_MLC1 | KAH0476563.1 | KAH0475263.1 |
| GAP40 | KAH0477413-4.1 | KAH0478356.1 |
| GAP45 (partial 3') | KAH0473219.1 | KAH0477289.1 |
| GAPM1 | GAPM3 KAH0485741.1<br>GAPMx KAH0485173.1<br>GAPMx KAH0472651.1 | GAPM3 KAH0485644.1<br>GAPMx KAH0481982.1 |
| GAC | KAH0484909-10.1 | KAH0477618:20.1 |
| ROM4 | KAH0485928-9.1<br>KAH0475712:14.1 | KAH0480431:33.1<br>KAH0472445-6.1 |
| AKMT | KAH0472731.1 | KAH0488385.1 |
| CDPK1(Tg)/CDPK4(Pf) | KAH0477425:27.1 | KAH0474276:78.1 |
| CDPK3(Tg)/CDPK1(Pf) | KAH0482406.1 | KAH0483722:23.1 |
| CDPK5(Pf)/CDPK5(Tg) | KAH0475451:53.1 | KAH0475693:95.1 |
| DGK1 | KAH0473632-3.1 | KAH0476753.1 |
| DOC2.1 | KAH0486912:14.1 | KAH0488625:27.1 |
| TSP-1 (a) | KAH0483741:43.1 | KAH0487684-5.1 |
| TSP-2 (a) | KAH0474072.1 | KAH0482614-5.1 |
| TSP2 (a) | KAH0472958:61.1 | KAH0473100:103.1<br>KAH0473117.1 |
| TSP_EGF-1 (a) | KAH0477971:73.1 | KAH0472910:12.1 |
| TSP_EGF-2 (a) | KAH0483270-1.1 | KAH0483538:40.1 |

(a) TRAP like candidates

**Table S8. *P. cf. gigantea* A and B glideosome and TRAP-like proteins identifiers**, related to Figure 6.
